# Conformational dynamics of cohesin/Scc2 loading complex are regulated by Smc3 acetylation and ATP binding

**DOI:** 10.1101/2022.12.04.519023

**Authors:** Aditi Kaushik, Thane Than, Naomi J Petela, Menelaos Voulgaris, Charlotte Percival, Peter Daniels, John B Rafferty, Kim A Nasmyth, Bin Hu

## Abstract

The ring-shaped cohesin complex is the key player in sister chromatid cohesion, DNA repair, and gene transcription. The loading of cohesin to chromosomes requires the loader Scc2 and is regulated by ATP. This process is also hindered by Smc3 acetylation. However, the molecular mechanism underlying this inhibition remains mysterious. Here we identify a novel configuration of Scc2 with pre-engaged cohesin and reveal dynamic conformations of the cohesin/Scc2 complex in the loading reaction. We demonstrate that Smc3 acetylation blocks the association of Scc2 with pre-engaged cohesin by impairing the interaction of Scc2 with Smc3’s head. Lastly, we show that ATP binding induces the cohesin/Scc2 complex to clamp DNA by promoting the interaction between Scc2 and Smc3 coiled coil. Our results illuminate a dynamic reconfiguration of the cohesin/Scc2 complex during loading and indicate how Smc3 acetylation and ATP regulate this process.

## Introduction

Sister chromatid cohesion is mediated by cohesin, a ring-shaped complex consisting of Smc1, Smc3 and the α-kleisin subunit Scc1. Smc1 and Smc3 are 50 nm long rod-shaped proteins with a “hinge” dimerization domain at one end and an ABC-like ATPase at the other. They dimerise through ‘hinge’ interaction and their ATPase heads are bridged by Scc1 to form a tripartite ring. The finding that Scc1’s cleavage by separase triggers sister chromatid disjunction led to the suggestion that cohesion is mediated by the co-entrapment of sister DNAs within a cohesin ring ^1^. Furthermore, cohesin has been demonstrated to act as a motor and regulate chromatin structure through loop extrusion ^2^.

To enable cohesin to entrap DNAs, one of three interfaces of cohesin must be opened during the loading reaction. Crucially, this process depends on the Scc2/4 loading complex, which is thought to catalyse ATP-dependent DNA entrapment (Murayama & Uhlmann 2014; Petela et al. 2018). However, the molecular detail of Scc2/4-mediated loading is still mysterious. The first step of loading is thought to involve Scc2 forming a transient complex with cohesin, called the cohesin/Scc2 loading complex. Subsequently, Scc2 stimulates ATP hydrolysis and opens the pre-assembled cohesin ring, permitting DNA entrapment (Hu et al. 2011; Murayama et al. 2018; Srinivasan et al. 2018). It is still disputed how the cohesin ring is opened. Genetic and biochemical evidence suggested that this reaction might trigger the opening of the Smc1/Smc3 hinge interface ^8,9^. Alternatively, recent studies proposed the Smc3/kleisin interface as the entry gate ^10,11^.

The loading process is more complicated than expected because cohesin exists in two different configurations. Cohesin coiled coil CCs are normally juxtaposed (J-cohesin), restricting head engagement (Fig S1A). The J-cohesin entraps DNA entrapment within the J-K compartment confined by kleisin and the head domains ^12^. ATP binding drives head engagement (E-cohesin) and the separation of head-end CCs, which are crucial for ATP hydrolysis and loading (Fig S1A). Recent Cryo-EM studies revealed that Scc2 can interact with both J- and E-cohesin ^11,13–15^, suggesting a dynamic reconfiguration of the Scc2/cohesin complex during the loading reaction (Fig S1A).

The release of DNAs from cohesin rings is achieved either through kleisin cleavage by separase, which occurs at the onset of anaphase ^16^, or through other stages of the cell cycle via a separase-independent mechanism ^17^. The latter requires a regulatory subunit Wapl ^18–21^. It induces the opening of the Smc3/kleisin interface in an ATP-dependent manner, thereby creating an exit gate permitting entrapped DNAs to escape from the cohesin ring ^22^. This Wapl-dependent release also detaches the native separase-cleaved N-Scc1 fragment from Smc3, which was used as an index of releasing activity ^23^.

Both Scc2- and Wapl-triggered ring openings could also pose an inherent risk to cohesion and, therefore, must be neutralized once sister chromatid cohesion is established. During DNA replication, Eco1, an acetyltransferase, transfers acetyl groups to Smc3 K112 and K113 in cohesin complexes destined to hold sister chromatids together ^18,24,25^. This modification is believed to antagonise Wapl-dependent releasing activity by preventing the opening of the Smc3/kleisin interface ^22^. Surprisingly, pre-acetylated cohesin fails to establish cohesion and a deacetylase, Hos1 in yeast or HDAC8 in humans, is required to recycle acetylated cohesin by removing acetyl groups ^26–28^. Compromising HDAC8 activity in humans leads to Cornelia de Lange syndrome (CdLS), a cohesin-related genetic disorder. A clue to how acetylated Smc3 hinders cohesion derives from a study of the acetylation mimicking form of Smc3, *smc3 K112Q K113Q* (referred to simply as *smc3QQ* hereinafter) ^29^. Similar to pre-acetylated cohesin, this version of Smc3 cannot create cohesion and support cell growth. Calibrated ChIP-seq revealed that *smc3QQ* significantly impairs cohesin loading *in vivo* ^29^. This implies that Smc3 acetylation not only counteracts Wapl-dependent releasing activity but also inhibits the loading reaction. It is not clear whether Smc3 acetylation blocks these opposite processes. One possibility is that this modification simply prevents the opening of the Smc3/kleisin interface because this opening is claimed to be required by both loading and release ^11,22^. Alternatively, Smc3 acetylation might regulate these processes via different mechanisms. To understand the role of Smc3 acetylation in loading and release, these possibilities must be clarified.

In this study, we investigated the mechanism by which Smc3 acetylation inhibits the loading reaction using the acetylation mimicking mutant *smc3QQ*. We first demonstrated that Smc3 acetylation impacts loading and release through independent pathways. To understand how Smc3 acetylation regulates loading, we established the dynamics of conformational changes in the Scc2/cohesin complex during loading by corroborating the two reported configurations with a novel transitional configuration identified in this study. We then showed that Smc3 acetylation inhibits the transitional configuration by blocking the interaction of Scc2 with the Smc3 head. Additionally, we revealed that ATP binding promotes Scc2/Smc3 CC interaction to regulate DNA clamp. In light of our results, we propose a dynamic reconfiguration of the Scc2/cohesin complex during loading regulated by Smc3 acetylation and ATP binding.

## Results

### Screening of mutations suppressing the lethality of *smc3QQ*

To understand the roles of Smc3 acetylation in loading and release, we searched for mutations that suppress the lethality of *smc3QQ*. Because loading, not release, is essential for sister chromatid cohesion and cell proliferation, these suppressor mutations should restore the loading of *smc3QQ*. If Smc3 acetylation inhibits loading and release through the same mechanism, namely preventing the opening of the Smc3/kleisin interface, these mutations should also improve the releasing activity of *smc3QQ*. In a screen for suppressors of synthetic lethality caused by *wpl1Δ* and *scc2-45*, we identified two suppressor mutations *smc3E199A* and *smc3R1008I* (Fig S1B). Though at opposite ends of the Smc3 polypeptide, E199 and R1008 are situated close to each other on opposite strands of Smc3’s anti-parallel CC, in a part that forms a three helical bundle with the α3 helix within Scc1’s NTD ^30^ (Fig. 1B). Interestingly, these mutations partially restored the proliferation of *smc3QQ* cells (Fig 1A). This suppressive effect was fairly weak as the growth of *smc3QQ R1008I* cells was poor at 25°C and stopped at 37°C (Fig S1C). In fact, *R1008I* barely suppressed the loading defect of ectopically expressed *smc3QQ* (Fig 1C). This might be due to the competition of endogenous Smc3 for loading. To enable further characterisation of the suppressing effect, we evolved *smc3QQ R1008I* by introducing random mutations across *smc3QQ R1008I* and identified 34 mutations that further improved the growth of *smc3QQ R1008I* cells. Strikingly, 18 suppressor mutations are distributed across both strands of Smc3 CC (from R248 to N517 and from K689 to L965) (Fig 1D). This reveals that the CC domain has a crucial role in regulating loading. Without exception, all of these mutations require the primary mutation *R1008I* to permit the proliferation of *smc3QQ* cells, implying a central role of the *R1008* region in loading (data not shown). In this study, we used suppressor mutations, *W483R R1008I*, to investigate the role of Smc3 acetylation in loading, because the growth of *SMC3* and *smc3QQ W483R R1008I* cells are indistinguishable (Fig 1E).

**Fig 1:**
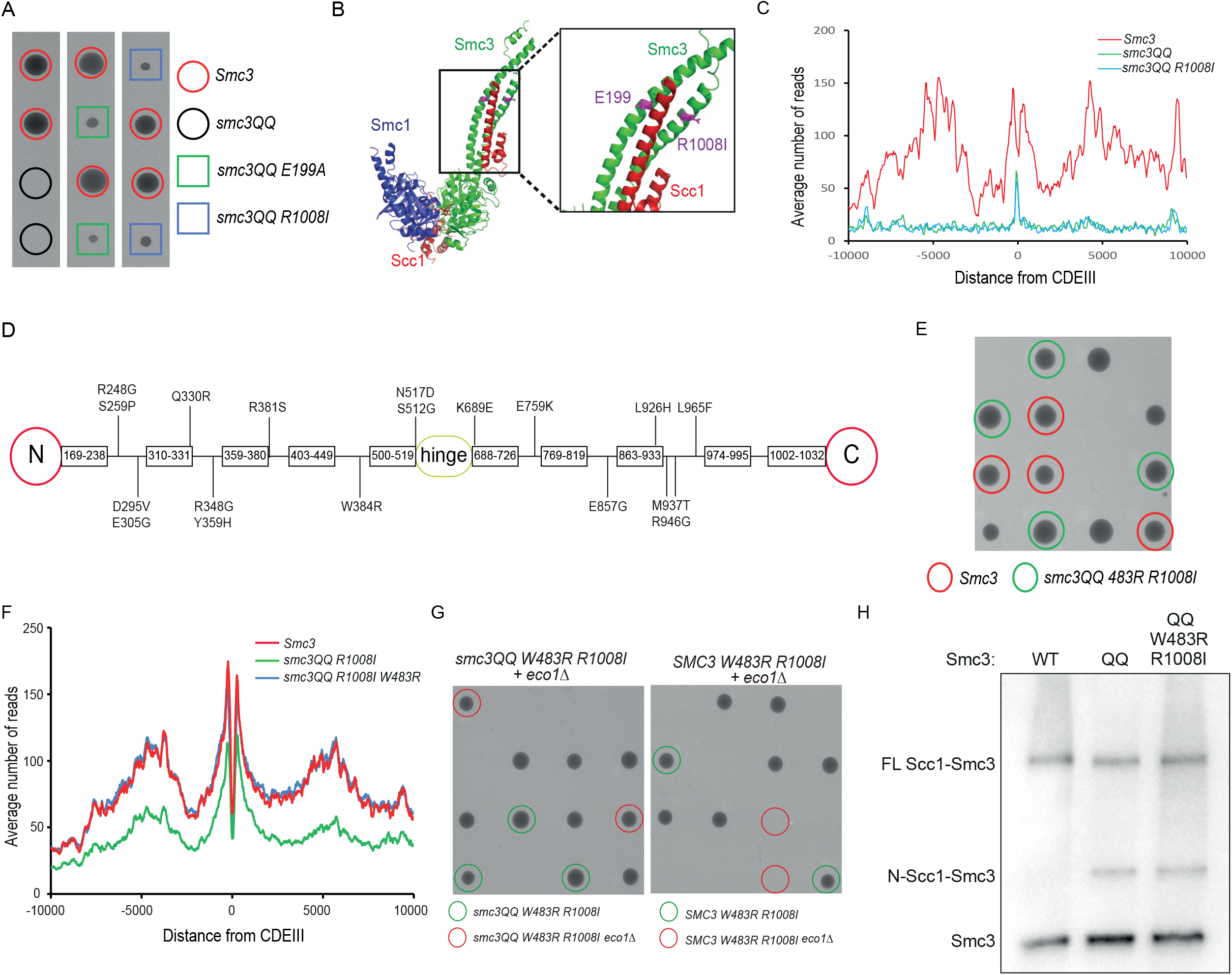
Smc3 acetylation affects loading and release through distinct mechanisms. (A) Tetrad dissection revealed Smc3 R008I and Smc3 E199A partially restored cell proliferation of *smc3 QQ* at 25°C. (B) Positions of smc3 E199 and R1008 on Smc3’s coiled coil. (C) Calibrated ChIP-seq revealed the defective DNA association of smc3QQ and smc3QQ R1008I (D) The distributions of second suppressor mutations on Smc3 CC (E) Restored growth by *smc3QQ* R1008I W483R. The growth of *Smc3* (red circles) and *smc3QQ W483R R1008I* (green circles) cells are indistinguishable. (F) Calibrated ChIP-seq profiles of Smc3, smc3QQ R1008I, and smc3QQ W483R R1008I. (G) Tetrad dissection revealed that smc3QQ W483R R1008I but not Smc3 W483R R1008I can bypass the requirement of Eco1 for cell growth. (H) Exponentially grown yeast cells were subject to in vivo BMOE crosslink. The free Smc3, Smc3s crosslinking to N-Scc1 and full-length Scc1 were analysed by western blot.

### Smc3 W483R R1008I fully suppresses *smc3QQ*’s defect in loading but not release

To measure the DNA association of Smc3QQ W483R R1008I without interference from endogenous Smc3, we replaced wild-type *SMC3* with *smc3QQ R1008I* or *smc3QQ W483R R1008I* and examined their DNA association using calibrated ChIP-seq. Smc3QQ R1008I showed modest DNA association, consistent with the ability of *smc3QQ R1008I* to weakly support cell growth (Fig 1F). Strikingly, *W483R R1008I* improved the loading of Smc3QQ to the level of wild-type Smc3.

We next addressed whether these suppressor mutations also restored the releasing activity of *smc3QQ*. During DNA replication, the releasing activity is inhibited by Eco1, which is critical for cohesion and cell viability ^18,24,25^. Therefore, the recovered releasing activity of *smc3QQ* by *W483R R1008I* will impair cohesion and lead to lethality in *eco1*Δ cells. Tetrad analysis showed that *smc3QQ W483R R1008I* bypassed the requirement of Eco1 for cell growth (Fig 1G). This effect is due to the acetylation-mimicking phenomenon of K112Q K113Q because Eco1 is still essential in *smc3 W483R R1008I cells* (Fig 1G). This genetic evidence demonstrated that *smc3QQ* can inhibit the releasing activity and the loading suppressor mutation *W483R R1008I* does not affect this inhibition.

To confirm this, we directly measured the releasing activity of *smc3QQ W483R R1008I* by examining the disassociation of the separase-cleaved N-terminal Scc1 fragment from Smc3 using *in vivo* BMOE crosslink between Smc3 S1043C and Scc1 C56 ^23^. As previously reported, the release of N-Scc1 from Smc3 is greatly impaired by s*mc3QQ*, and this defect cannot be remedied by *W483R R1008I* (Fig 1H).

All these results revealed that *smc3QQ* sufficiently prevents Wapl-mediated release and maintains cohesion if its loading defect is suppressed, demonstrating that Smc3QQ is not just a structural analogue of acetylation but also mimics its function. This allows us to use *smc3QQ* to investigate the biochemical properties of Smc3 acetylation. Moreover, *W483R R1008I* can fully suppress the defect of Smc3QQ loading, but not release, revealing that Smc3 acetylation prevents loading and release through different mechanisms. Hereby, we focused on the role of this modification in loading by studying *smc3QQ*.

### *smc3QQ* inhibits Scc2-stimulated ATP hydrolysis of cohesin

To understand how Smc3 acetylation inhibits loading, we first evaluated the effect of *smc3QQ* on the Scc2-stimulating ATPase activity of cohesin. In this assay, a recombinant tetramer of cohesin (Smc1, Smc3 or smc3QQ, Scc1, Scc3) was incubated with ATP and GFP-Scc2 ^4^. Consistent with previous reports, ATPase activity of the wild-type tetramer was detected only upon stimulation by Scc2 and DNA (Fig 2A) (Murayama & Uhlmann 2014; Petela et al. 2018). However, Scc2’s ability as a stimulator of ATPase activity was significantly undermined by *smc3QQ* (Fig 2B). This suggests that Smc3 acetylation and its mimicking QQ mutation impair the intrinsic ATPase activity of cohesin or inhibit the Scc2/cohesin interaction required for triggering ATP hydrolysis. Surprisingly, DNA almost fully recovered the Scc2-dependent ATP activity of *smc3QQ* (Fig 2B). This result is difficult to reconcile with the notion that *Smc3 K112 K113* acts as a DNA sensor to promote the DNA/cohesin association ^10^. In contrast, this finding reveals that DNA enhances Scc2- triggered ATP hydrolysis of Smc3QQ up to the wild-type level, indicating that DNA might actually facilitate ATP hydrolysis-compatible conformation of cohesin, for example, head engagement, or stabilise cohesin/Scc2 interaction, even when Smc3 is acetylated. However, Smc3QQ cannot be recruited to DNA *in vivo*, therefore, the effect of DNA cannot be achieved. Both loading and ATP hydrolysis require Scc2/cohesin interaction and ATP-dependent head engagement (Hu et al. 2011; Petela et al. 2018), thus, *smc3QQ* may affect one of these processes.

**Fig 2:**
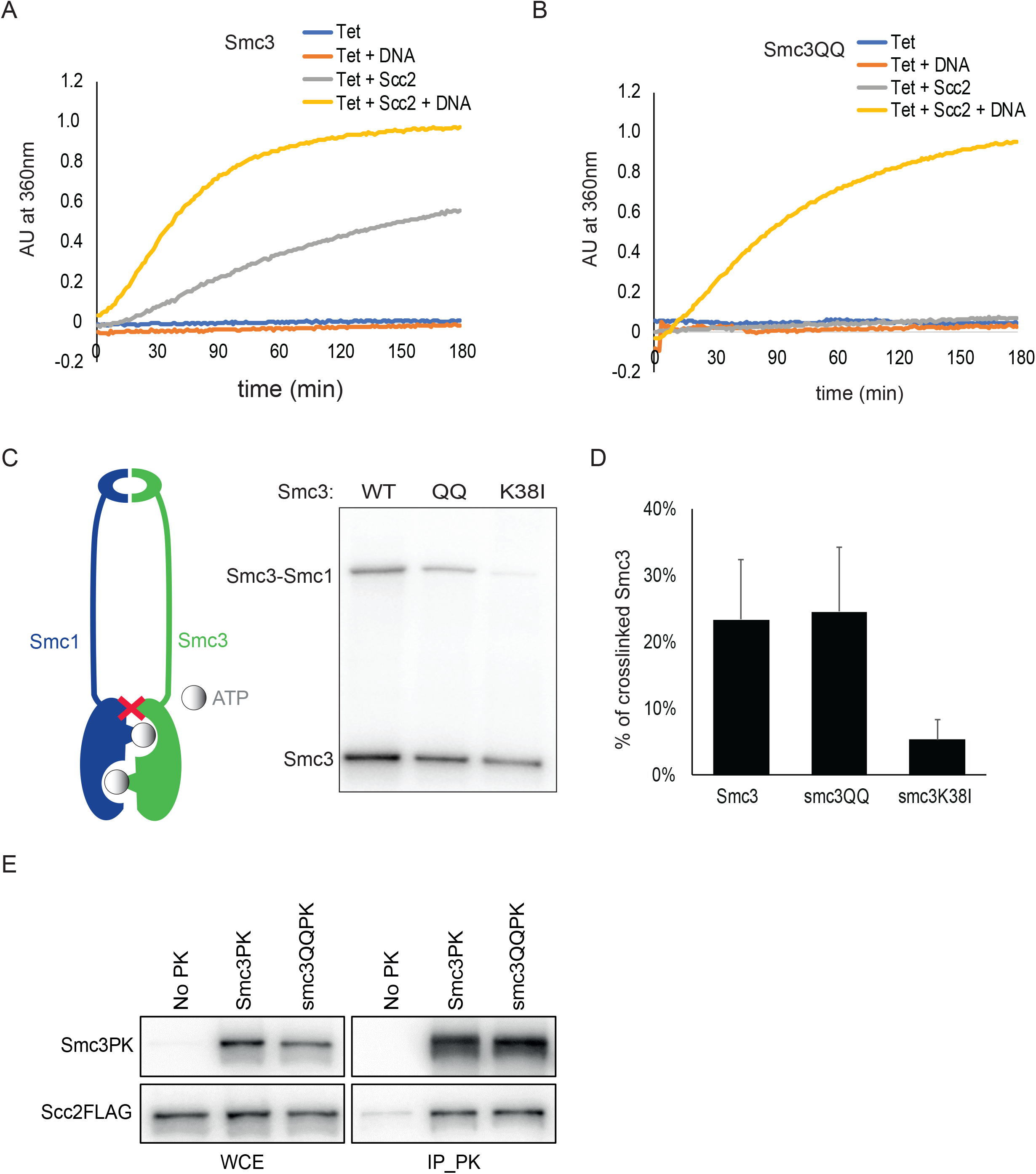
Smc3QQ affects cohesin’s interaction with Scc2 rather than DNA. (A) Recombinant tetramer of cohesin (Smc1, Smc3, Scc1 and Scc3) was incubated with DNA or Scc2 or both. ATP was added to initiate the reaction, and the reaction rate was measured as the change in absorption at 360nm over time. (B) ATPase activity of smc3QQ in the presence of Scc2 or DNA or both. (C) *In vivo* BMOE crosslink between Smc1 N1192C and Smc3 R1222C measure head engagement between Smc1 and Smc3, smc3QQ or smc3K38I. Crosslink bands were separated by gradient gel and Smc3 was examined by Western Blot. (D) The percentage of crosslinking efficiency in panel C was calculated as mean+SD of 3 independent experiments. WT crosslinking efficiency was 23%, Smc3QQ was 24% and Smc3K38I was reduced to 5%. (E) Co-immunoprecipitation of Scc2 with Smc3 or smc3 QQ.

### *smc3QQ* has little effect on head engagement and the assembly of Scc2/cohesin complex

It was proposed that Smc3 acetylation might inhibit head engagement because acetylated residues K112 and K113 are adjacent to the ATP binding pocket and this modification might alter its conformation ^30^. To survey this, we examined head engagement using the cysteine pair, Smc1 N1192C and Smc3 R1222C, which can be crosslinked by bismaleimidoethane (BMOE) only if the heads are engaged ^12^. Approximately 23% of Smc3 R1222C formed a covalent crosslink product with Smc1 N1192C with BMOE (Fig 2C and 2D). This crosslink occurs in an ATP-dependent manner because the ATP-binding mutant *smc3K38I* significantly reduced the crosslinking efficiency (5%) (Fig 2C and 2D). A similar conclusion was drawn by measuring the crosslinking efficiency of Smc1N1192C (Fig S2A and S2B). Interestingly, *smc3QQ* has little effect on the crosslink, with a crosslink efficiency of 24%. This result excluded the possibility that the defective loading of acetylated Smc3 is due to impaired head engagement.

Next, we examined whether Smc3 acetylation might prevent the Scc2/cohesin interaction. Co-immunoprecipitation revealed that Smc3QQ displayed a comparable ability to interact with Scc2 as wild-type Smc3 (Fig 2E). This indicates that the global interaction between Scc2 and cohesin was barely impaired by Smc3 acetylation.

### Scc2 interacts with both Smc1 and Smc3 head domains

Although *smc3QQ* has no discernible effect on the assembly of the Scc2/cohesin complex, the possibility of this mutant altering a local contact and disrupting an essential Scc2/cohesion configuration cannot be ruled out. Based on this rationale, we performed an *in vivo* BPA crosslink screen to identify interfaces between cohesin and Scc2. Considering its role in ATP hydrolysis, Scc2 is likely to interact with cohesin heads. We thus reasoned that these three regions of the Smc3 head are potential interaction sites. The first region is the Smc3 acetylation site, a highly conserved KKD strand. The second candidate site is the adjacent α helix (R58-L64), presumably serving as a communicator between the KKD strand and the Smc3 ATP-binding pocket ^30^. Lastly, we inferred that the regions proximal to the Smc3 ATP-binding pocket could also be potential sites for interaction with Scc2. To survey this, we expressed a set of Smc3 variants in which a given residue (K57, R58, R61, M74, H67, Q67, L111, or Q117) was substituted with p-benzoyl-l-phenylalanine (BPA) (Fig 3A), a photoreactive amino acid that can crosslink to any other residues within a distance of 7Å upon UV irradiation ^33,34^. A plasmid-shuffling assay demonstrated that these Smc3 variants can replace wild-type Smc3 to maintain cell viability (data not shown). We did not detect any Smc3-Scc2 crosslink at the KKD strand (L111BPA and Q117BPA), suggesting that Scc2 might not directly interact with the Smc3 acetylation site (Fig 3B). Interestingly, we found that Scc2 was covalently crosslinked with Smc3 K57BPA, Q67BPA, and M74BPA, among which Smc3 Q67BPA showed the highest crosslinking efficiency (Fig 3B). In fact, Smc3 Q67BPA formed two distinct crosslink products (Fig S3A). Increasing the size of Scc2 by fusing it with a 6xFlag tag led to an upshift of the top crosslink band, indicating an Smc3-Scc2 crosslink (Fig S3A). This was confirmed by Western blot using an anti-FLAG antibody. This result reveals a physical interaction between Scc2 and the Smc3 ATP-binding domain since Smc3 Q67 is located on the boundary of Smc3’s ATP-binding pocket (Fig 3A). Fusion of Smc1 with a 9xMyc tag revealed that Smc3 Q67BPA was also crosslinked to Smc1 (Fig S3B). This was not surprising, given that head engagement brings Smc3 Q67 close to Smc1’s head domain (Fig 3A).

**Fig 3:**
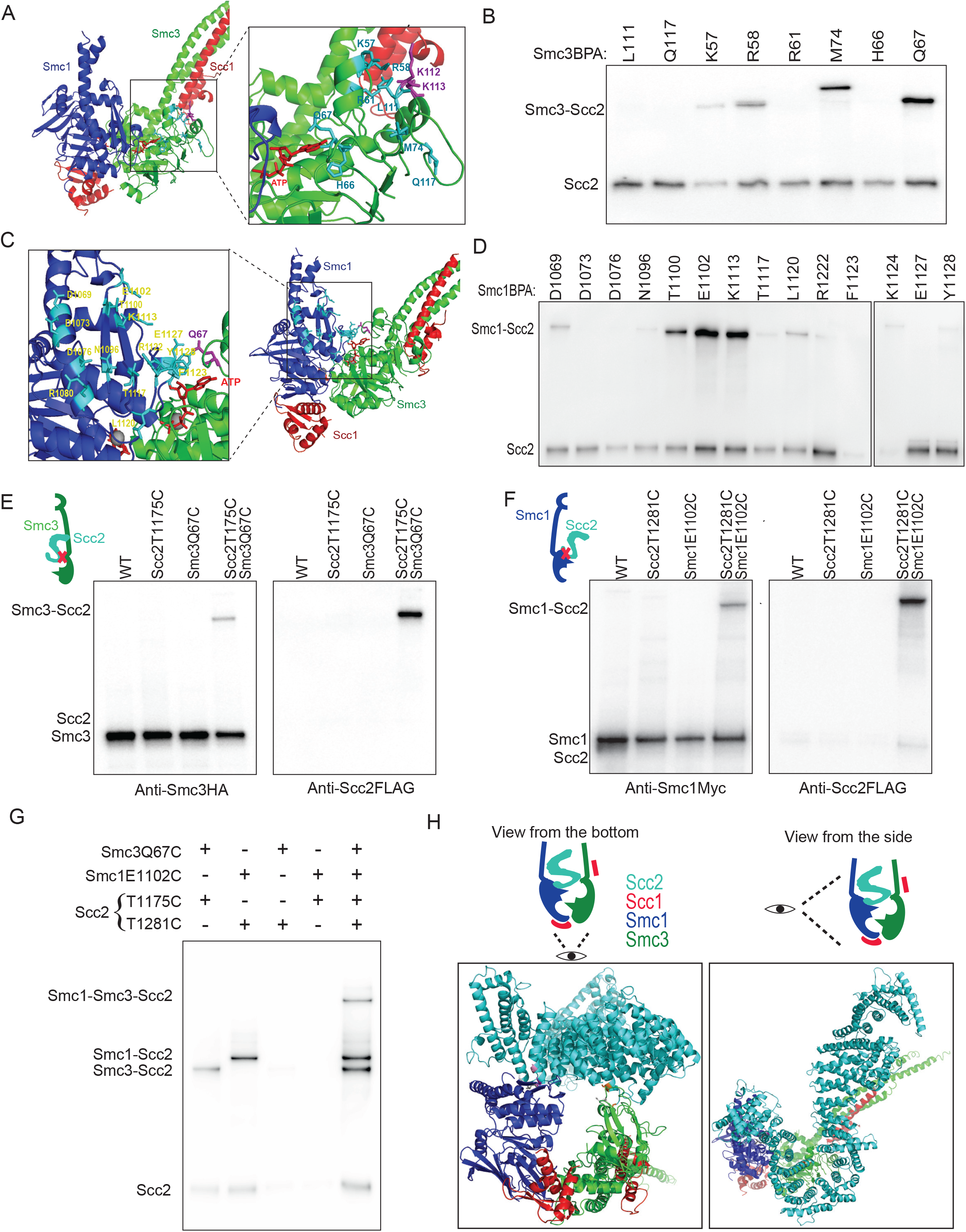
Mapping of Smc1-Scc2 and Smc3-Scc2 interfaces. (A) The indicated residues near the Smc3 KKD strand and ATP binding pocket were substituted by BPA. B) The residues of Smc3 were selected to perform *in vivo* BPA crosslinking. After immunoprecipitated against Scc1-PK and separated on gradient gel, Scc2 was analysed by western blot. Smc3 K57BPA, Q67BPA and M74BPA successful crosslinked with Scc2, in which Smc3 Q67BPA showed the highest crosslink efficiency. (C) The indicated residues around the Smc1 ATP binding pocket were selected to perform *in vivo* BPA crosslink. (D) After UV crosslink, cohesin complex was immunoprecipitated against Scc1-PK and Scc2 was detected by Western Blot. Smc1 E1102BPA and Smc1 K1113BPA showed higher crosslink efficiency. (E) *In vivo* crosslinking revealed close proximity between Smc3 Q67 and Scc2 T1175. Single or both residues were replaced by cysteine and *in vivo* BMOE crosslinking was performed, immunoprecipitated on Scc1-PK, and Smc3 or Scc2 were examined by western blot. Successful crosslink product formation depended on the presence of both cysteines. (F) *In vivo* crosslinking revealed close proximity between Smc1 E1102 and Scc2 T1281. (G) Smc3 Q67C/Scc2 T1175C and Smc1 E1102C/Scc2 T1281C were simultaneously crosslinked *in vivo* using BMOE. Immunoprecipitation was performed against Scc1-PK, separated on gradient gel, and Scc2 was analysed by Western Blot. (H) Structural model of a novel Scc2cohesin configuration.

We also carried out a BPA screen in a region of Smc1 that is part of the ATP-binding pocket (Fig 3C) and identified three adjacent BPA substitutions, Smc1 T1100BPA, E1102BPA, and K1113BPA, which can crosslink to Scc2 (Fig 3D). Like Smc3 Q67BPA, Smc1 E1102BPA also crosslink to Smc3 (Fig S3C). To our surprise, Smc1-Scc2 crosslinks were barely observed for Smc1 F1123BPA, E1127BPA, and Y1128BPA. Since head engagement brings these residues close to Smc3 Q67 and their side chains point in the same direction (Fig 3C), if Scc2 interacts with Smc3 Q67 with engaged heads, these three BPA-substituted Smc1 variants should also crosslink with Scc2. Hence, this result suggests that heads are not engaged when Scc2 interacts with Smc3 Q67. Through these experiments, we identified two interfaces between Scc2 and the cohesin heads.

### Both interfaces co-exist in an Scc2/cohesin configuration

To define the interface between Scc2 and Smc3 Q67, we developed a novel strategy to map Scc2 sites crosslinked by Smc3 Q67BPA. To do this, we created a set of functional Scc2 alleles, each containing 3x TEV protease cleavage sequences at the indicated sites (Fig S3D). All these versions of Scc2 are labelled with a 6xFLAG tag at the C-terminus, except Scc2TEV1176, because the FLAG-fusion destroys its function. To determine the region of Scc2 crosslinked by Smc3-Q67BPA, the Smc3/Scc2 crosslink products were subjected to TEV protease digestion, producing an untagged Scc2 N-terminal fragment and a FLAG-tagged C-terminal fragment (Fig S3D). Western blot showed that Smc3 Q67BPA crosslinked to the C-terminal fragments when cleavage occurred in the region 215-1109, while it crosslinked to the N-terminal fragments when cleaved at positions 1176 and 1222 (Fig S3D). This revealed that the Scc2 residue crosslinked by Smc3 Q67BPA is located in the region of 1110-1176.

To further refine this interface, we replaced 15 residues across this region and Smc3 Q67 with cysteine (Fig S3E). Interestingly, six cysteine-substituted alleles of Scc2 (R1115C, N1133C, E1158C, D1162C, E1168C, and R1173C) are synthetically lethal with *smc3 Q67C*, implying an essential role of this interface in loading (data not shown). The proximity between the remaining nine cysteines and Smc3 Q67 was determined using *in vivo* BMOE crosslinking. We found the crosslinks of Smc3 Q67C with Scc2 S1171C or T1175C showed the highest efficiency, suggesting an interaction between Smc3 Q67 and Scc2 S1171/T1175 (Fig S3E and 3E). Using the same strategy, we found that Smc1 E1102BPA crosslinked to an Scc2 region between residues 1260-1303 (Fig S3F) and a further cysteine pair screen showed that Smc1 E1102C crosslinked to Scc2 T1281C with the highest efficiency, indicating the interface between Smc1 and Scc2 (Fig 3F and S3G).

To test whether both interfaces would exist in one Scc2/cohesin configuration or belong to two different configurations, we introduced both cysteine pairs into endogenous Smc1/Smc3/Scc2 and examined whether these two crosslinks could occur simultaneously. Indeed, the Scc2/cohesin complex bearing both cysteine pairs generated not only two single crosslink products but also an Smc1-Smc3-Scc2 double crosslink product (Fig 3G). The crosslink strictly took place between the cysteine pair of Smc3 Q67C/Scc2 T1175C or Smc1 E1102C/Scc2 T1181C since no crosslink product between any other cysteine pair combination was observed (Fig 3G). The double crosslink demonstrates that the two interfaces (Smc3 Q67C/Scc2 T1175C and Smc1 E1102C/Scc2 T1181C) co-exist in an Scc2/cohesin configuration. Based on the proximity of these two cysteine pairs, we constructed a structural model of this configuration (Fig 3G).

### Smc3 Q67/Scc2 interface represents a novel configuration of the Scc2/cohesin complex

In both reported configurations of Scc2/J-cohesin ^15^ and Scc2/E-cohesin ^11,13,14^, the predicted distances between Smc3 Q67 and Scc2 T1175 are more than 16nm, beyond the BMOE-crosslink limit (Fig S4A and S4B), suggesting that Smc3 Q67/Scc2 interface might present in a novel configuration of the cohesin/Scc2 complex. However, we cannot exclude the possibility that the flexible Smc3 Q67 loop can swing to Scc2 T1175 in the Scc2/E-cohesin complex. Interestingly, a neighbour residue Smc3 M74 is found to be in the vicinity of Scc2 in our modelled Scc2/cohesin and also in Scc2/E-cohesin configurations, consistent with our result that Smc3 M74BPA can crosslink to Scc2 (Fig 3B). However, Smc3 M74 is predicted next to different Scc2 residues, Scc2 N886 in our modelled structure and E821 in Scc2/E-cohesin configuration (Fig S4C and S4D), which is confirmed by BMOE crosslink (Fig 4B). If these crosslinks take place only in the Scc2/E-cohesin configurations due to the flexibility of the Q67/M74 loop, allowing Smc3 M74C to oscillate between Scc2 N886 and E821, this movement will also bring Smc3 M74 close to Scc2 R830 (Fig 4A). This possibility is excluded as no crosslinking was observed between this cysteine pair (Fig 4B), which supports our claim of a novel Scc2/cohesin configuration indicated by the Smc3 Q67C/Scc2 T1175C crosslink.

**Fig 4:**
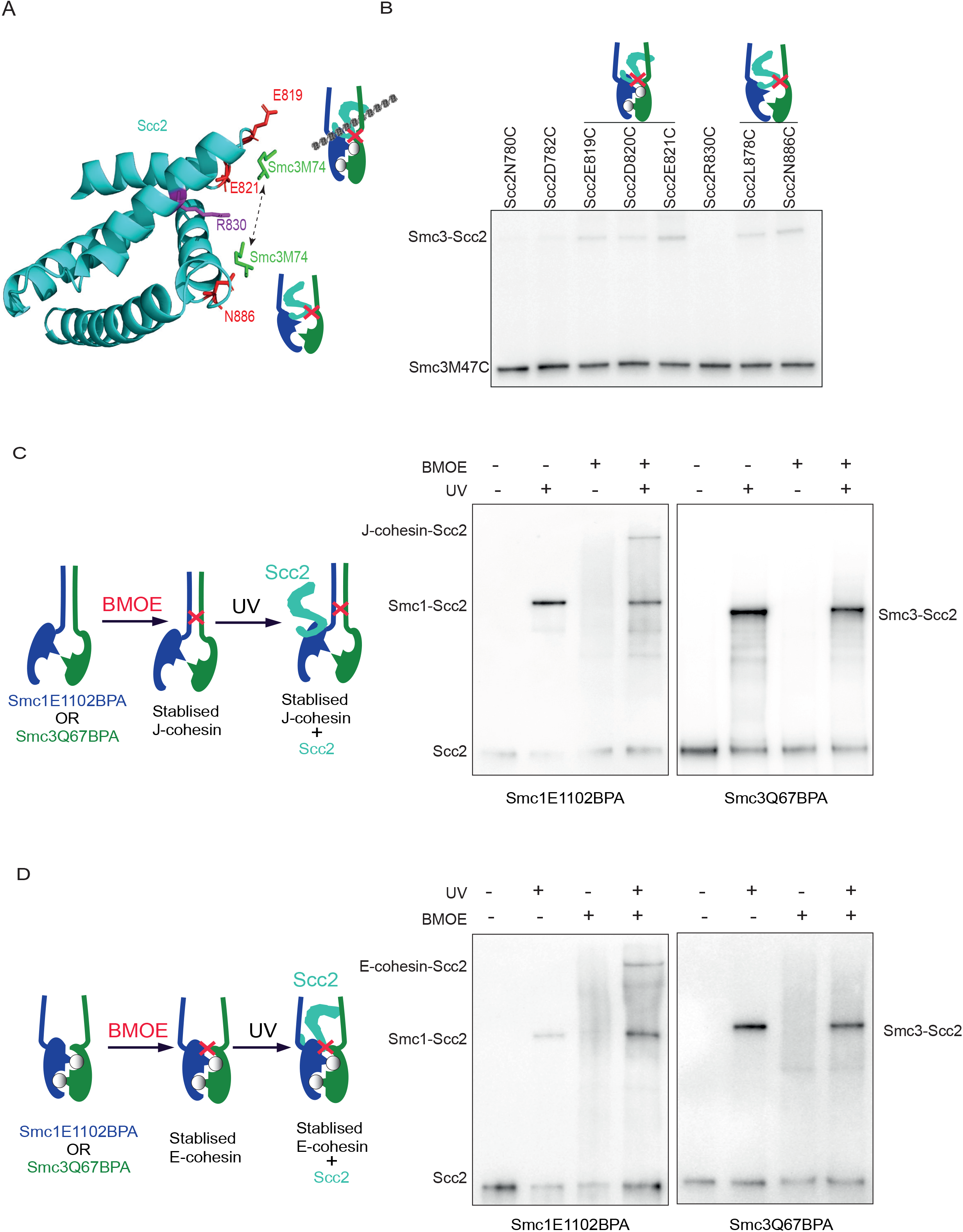
Smc3 Q67/Scc2 T1175 interface represents a novel Scc2/cohesin conformation. (A) The related positions of Smc3 M74 with Scc2 E819, E820, R830, and N886 in the hypothetical oscillation model. (B) The BMOE crosslink of Smc3 M74C with indicated cysteine substituted Scc2. Scc2 was detected by Western Blot. (C) *In vivo* BPA crosslink of Smc1 E1102BPA or Smc3 Q67BPA to Scc2 in J-cohesin. Juxtaposed CCs were stabilised by *in vivo* BMOE crosslinking Smc1 R1031C and Smc3 E202C, followed by UV-induced BPA crosslink. Immunoprecipitated on Scc1-PK, separated on gradient gel, and Scc2 was analysed by Western Blot. (D) *In vivo* BPA crosslink of Smc1 E1102BPA or Smc3 Q67BPA to Scc2 in E-cohesin. Engaged heads were stabilised by *in vivo* BMOE crosslinking of Smc1 R1031C and Smc3 E202C, followed by UV-induced BPA crosslink. Immunoprecipitated on Scc1-PK, separated on gradient gel, and Scc2 was analysed by Western Blot. (E) Summary of structural features of three different Scc2/cohesin complexes.

The structural model of the new configuration predicts two facts: separated cohesin CCs, at least from neck to head, and disengaged heads. This indicates that the cohesin complex in this configuration is neither J-cohesin nor E-cohesin. By this rationale, Smc3 Q67BPA from either J- cohesin or E-cohesin should not crosslink to Scc2. To test this, we first chemically stabilised J- cohesin or E-cohesin by crosslinking the juxtaposed CCs or engaged heads using BMOE. We then examined whether Smc3 Q67BPA can crosslink to Scc2 (Fig 4C and 4D, left panels). If BMOE-stabilised J-cohesin or E-cohesin still interacts with Scc2, the following BPA crosslink would produce a triple crosslink product among Smc1-Smc3-Scc2. Otherwise, the BMOE crosslink of Smc1/Smc3 and BPA-crosslink of Smc3 Q67BPA/Scc2 should be mutually exclusive. In this experiment, Smc1 E1102BPA was used as a positive control, as the Cryo-EM structures revealed that the Smc1 E1102/Scc2 interface presents in both Scc2/J-cohesin and Scc2/E-cohesin configurations (Petela et al. 2021; Shi et al. 2020; Murayama & Uhlmann 2020; Collier et al. 2020). To identify the cysteine pair for the crosslink of the juxtaposed CCs, we constructed a structural model of the juxtaposed CCs (Fig S4F) and found a cysteine pair (Smc1 R1031C/Smc3 E202C) that can be used to covalently fix the J-cohesin (Fig S4G). As predicted, Smc1 E1102BPA, but not Smc3 Q67BPA, was able to crosslink with Scc2 when the juxtaposed CCs were stabilised by a BMOE crosslink (Fig 4C). We repeated this experiment with engaged heads crosslinked using a cysteine pair (Smc1 N1192C and Smc3 R1222C) ^12^. Unlike Smc1 E1102BPA, Smc3 Q67BPA did not crosslink with Scc2 when heads were engaged (Fig 4D). All these results show that the cohesin/Scc2 configuration characterised by the Smc3 Q67/Scc2 interface is distinct from the Scc2/J-cohesin or Scc2/E-cohesin configurations, indicating that Smc3 Q67/Scc2 interface represents a novel configuration.

This configuration shares some features with these two reported configurations: Scc2/J-cohesin and Scc2/E-cohesin (Fig 4E). Like in the Scc2/J-cohesin configuration, cohesin heads are disengaged in the new configuration. However, cohesin CCs are apart and Scc2 can interact with both heads, as in the Scc2/E-cohesin configuration. These features of the new configuration suggest it might be an intermediate configuration in the transition from Scc2/J-cohesin to Scc2/E-cohesin/DNA. Therefore, we named this novel configuration Scc2/pre-E-cohesin.

### *smc3QQ* impairs the formation of Scc2/pre-E-cohesin and Scc2/E-cohesin configurations, but not Scc2/J-cohesin

Identification of the Scc2/pre-E-cohesin configurations, together with two reported configurations, allows us to elucidate the effect of Smc3 acetylation on loading by investigating how *smc3QQ* affect these configurations. We first surveyed the effect of *smc3QQ* on Scc2/pre- E-cohesin configuration by examining the Smc3 Q67C/Scc2 T1175C crosslink. To avoid the interference caused by Smc3 acetylation, cells were arrested in the late G1 phase by overexpression of Sic1(9m), a nondegradable Sic1 (Chan et al., 2012). At this stage, Smc3 is not acetylated because this modification depends on DNA replication. The crosslink between Smc3 Q67C and Scc2 T1175C is dramatically reduced by *smc3QQ*, to about 10% of wild-type Smc3 (Fig 5A, 5B and S5A). However, *smc3QQ* might only distort this interface and lead to the misalignment of these two residues. If it is the case, Smc3 Q67 might still interact with other Scc2 residues. To examine this possibility, we re-evaluated the effect of *smc3QQ* on Smc3 Q67BPA/Scc2 crosslinks. The acetylation-mimicking mutant scarcely affects the crosslink of Smc3 Q67BPA with Smc1 (Fig S5B). This reconciles with the notion that Smc3 acetylation has little effect on head engagement (Fig 2D). However, the Smc3QQ Q67BPA/Scc2 crosslink product was barely detectable (Fig S5B). All these results revealed that the acetylation-mimicking mutant seriously impairs the Scc2/pre-E-cohesin configuration The Scc2/E-cohesin/DNA configuration reflects a key step in loading. If the Scc2/pre-E-cohesin configuration occurs prior to the Scc2/E-cohesin configuration, we inferred that *smc3QQ* would also negatively affect the Scc2/E-cohesin configuration. As revealed by Cryo-EM analysis, this configuration is maintained by three interfaces, including Smc1 E1102/Scc2 T1281. The other two interfaces, Scc2/Smc3 head and Scc2/Smc3 CC, seem unique to this configuration and form a compartment for DNA entrapment (Fig 5C). To detect these interfaces *in vivo*, we introduced two cysteine pairs into these interfaces, respectively, with a distance allowing BMOE crosslink (Fig 5C). As shown in Fig 5D and S5C, Smc3 head/Scc2 can be specifically crosslinked by BMOE *in vivo* with cysteine pair, Smc3 S72C and Scc2 E819C. Therefore, this crosslink can detect the *in vivo* interaction of the Scc2/Smc3 head presented in the Scc2/E-cohesin configuration. By the same principle, the crosslink relying on the cysteine pair of Smc3 K1004C and Scc2 D369C verifies the interface between Scc2 and Smc3 CC *in vivo* (Fig 5E and S5D). With these cysteine pairs, we found that *smc3QQ* dramatically decreased the crosslinks in both interfaces by about 90%, compared to the wild type (Fig 5F-5I, S5E, and S5F). These results resemble the inhibition of the crosslink between Smc3 Q67C and Scc2 T1175C caused by *smc3QQ*, to a similar extent. It suggests that *smc3QQ* impairs both the Scc2/pre-E-cohesin and Scc2/E-cohesin configurations.

**Fig 5:**
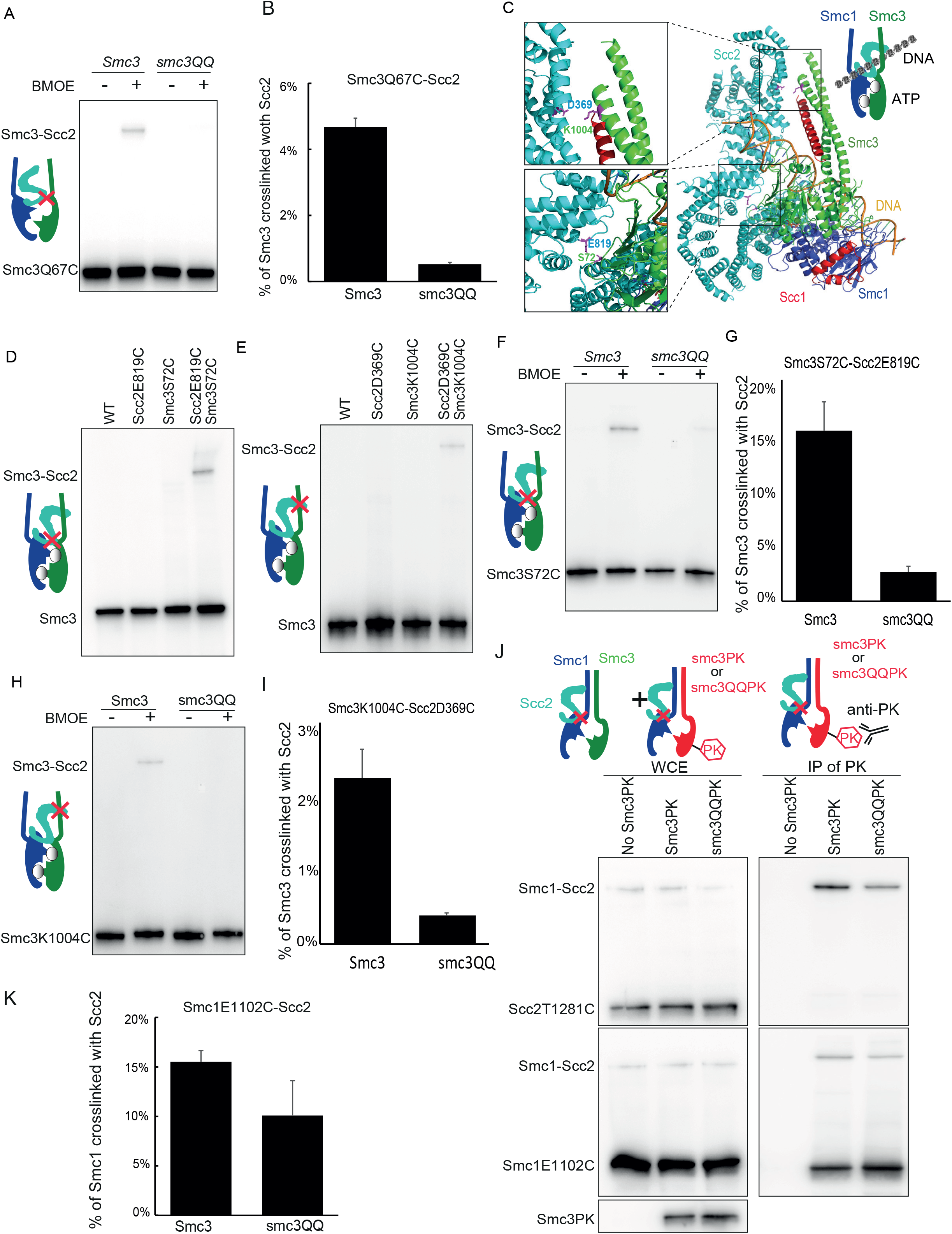
Inhibition of Scc2/pre-E-cohesin and Scc2/pre-E-cohesin configurations by *smc3QQ*. (A) *In vivo* BMOE crosslink of Scc2 T1175C with Smc3 or smc3QQ Q67C. Immunoprecipitated on Scc1-PK, separated on gradient gel and Smc3 was analysed by western blot. (B) The percentage of crosslinking efficiency in panel A was calculated as mean ± SD of 3 independent experiments. smc3QQ reduced the crosslinking efficiency to about 10% of wild type. (C) Indicated residues on the interface of Smc3 head/Scc2 or Smc3 CC/Scc2 in Scc2/E-cohesin configuration. (D and E) *In vivo* BMOE crosslinking between Smc3 S72C-Scc2 E819C (D) and Smc3 K1004C- Scc2 D369C (E) was performed using. Immunoprecipitated on Scc1-PK, separated on gradient gel and analysed by western blot. Successful crosslink product formation depended on the presence of both cysteines of each pair. (F) *In vivo* BMOE crosslinking of Scc2 E819C with Smc3 or smc3QQ S72C. (G) The percentage of crosslinking efficiency in panel F was calculated as mean±SD of 3 independent experiments. *smc3QQ* reduced the crosslinking efficiency to about 10% of wild type. (H) *In vivo* BMOE crosslinking of Scc2 D369C with Smc3 or smc3QQ K1004C. (I) The percentage of crosslinking efficiency in panel H was calculated as mean±SD of 3 independent experiments. *smc3QQ* reduced the crosslinking efficiency to about 10% of wild type. (J) Using BMOE, *in vivo* cysteine crosslink was performed between Scc2 T281C and Smc1 E1102C in cells with ectopically expressed Smc3-PK or smc3QQ-PK. Immunoprecipitation was performed using anti-PK antibody and the crosslinked Smc1-Scc2 was detected by Western blot. (K) The percentage of crosslinking efficiency in panel J was calculated as the mean±SD of 3 independent experiments. smc3QQ had a mild effect on the crosslink.

Even though *smc3QQ* dramatically impairs the Scc2/pre-E-cohesin and Scc2/E-cohesin configurations, Smc3QQ is still able to interact with Scc2 (Fig 2E). We deduced that the Scc2/J- cohesin configuration might be not affected by *smc3QQ*. However, we cannot identify a cysteine pair specific to this configuration because the only Scc2/cohesin interface (Smc1 E1102/Scc2 T1285) is also present in the other two configurations. If *smc3QQ* would have a limited effect on the crosslink of Smc1 E1102C/Scc2 T1285C, according to the process of elimination, the Scc2/J- cohesin configuration is barely affected by *smc3QQ*. To test this, we ectopically expressed PK-tagged Smc3 or smc3QQ and the cysteine pair Smc1 E1102C/Scc2 T1285C was introduced into endogenous Smc1 and Scc2. Therefore, its crosslink will occur in the cohesin complex with endogenous Smc3 or with ectopically expressed Smc3-PK/Smc3QQ-PK (Fig 5J). To specifically measure the effect of *smc3QQ* on the Smc1 E1102/Scc2 T1281 interface, the Smc1/Scc2/Smc3QQ-PK complex was first immunoprecipitated (Fig 5J). In contrast to Smc3 Q67C or S72C/Scc2 crosslink, the Smc1 E1102C/Scc2 crosslink was mildly affected by *smc3QQ* (Fig 5J and 5K). This conclusion was also supported by the Smc1 E1102BPA crosslink (data not shown). Altogether, these results demonstrated that Smc3 acetylation blocks loading by preventing the Scc2/pre-E-cohesin and Scc2/E-cohesin configurations and has little effect on the Scc2/J-cohesin configuration.

### smc3 W483R R1008I improves the interaction between Scc2 and smc3QQ’s head and CC

We have demonstrated that the suppressor mutation *W483R R1008I* restores the loading of Smc3QQ, thus this mutation probably permits the formation of these configurations impaired by *smc3QQ*. To examine this, we surveyed the effect of *W483R R1008I* on the crosslink between Smc3 Q67C and Scc2 T1175C. We found that *R1008I* mildly improved this crosslink (about 40% of wild-type Smc3 with Scc2), but *W483R* has little effect (Fig 6A, 6B and S6A). The combination of *R1008I* and *W483R* greatly increased the crosslinking efficiency, about 70% of the wild type. This effect was confirmed using smc3 Q67BPA/Scc2 crosslink (Fig S6B). Similar effects of these suppressor mutations were also found on the Smc3 S72C/Scc2 E819C interfaces (Fig S6D and S6E). It suggested that modification of the configuration of Smc3 CC conferred by *W483R R1008I* might affect the orientation of Smc3QQ’s head domain, which facilitates its interaction with Scc2. Interestingly, *R1008I*, regardless of *W483R*, improved Scc2/Smc3 CC interaction represented by Smc3 K1004C/Scc2 D369C crosslinks, about 70% of the wild type (Fig 6C, 6D and S6C). This suggests that another role of *R1008I* is to restore the interaction of Scc2 with Smc3 CC. R1008’s proximity to the Scc2/Smc3 CC interface supports this conclusion. These results revealed that the suppressor mutation *W483R R1008I* restored the configurations of Scc2 with pre-E- cohesin or E-cohesin which are impaired by *smc3QQ*.

**Fig 6:**
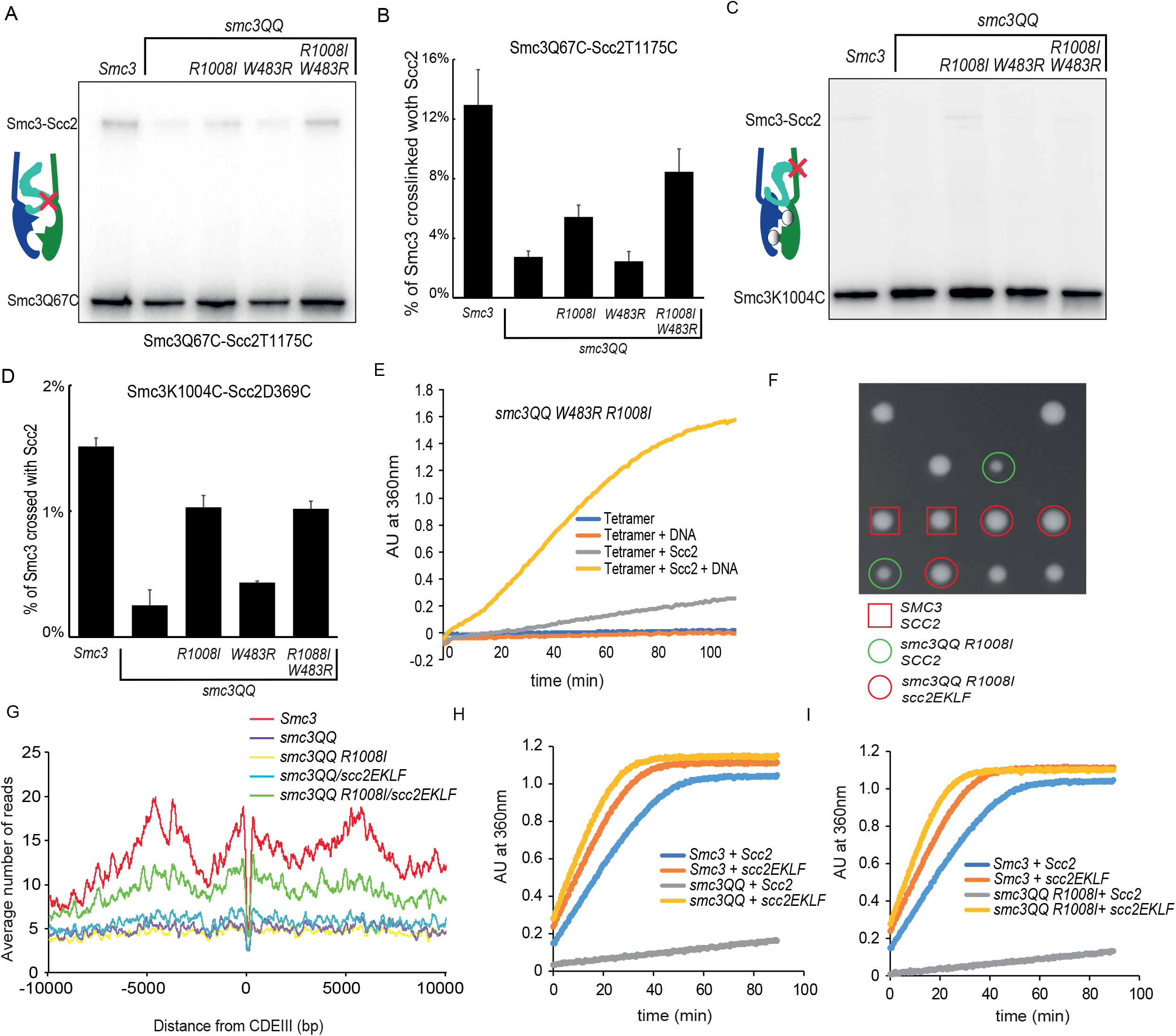
Loading defect of smc3QQ is rescued by Smc3 suppressors and a hypermorphic Scc2 allele. (A) Recombinant tetramer of cohesin smc3QQ W483R R1008I, Smc1, Scc1 and Scc3 was incubated with DNA or Scc2 or both. ATP was added to initiate the reaction and reaction rate was measured as the change in absorption at 360nm over time. (B) Using BMOE, *in vivo* cysteine crosslink was performed between Scc2 T1175C and smc3QQ Q67C in the presence of Smc3 R1008I or W483R or both. The first lane shows the Smc3 Q67C- Scc2 T1175C crosslink. Immunoprecipitated on Scc1-PK, separated on gradient gel, and analysed by western blot. (C) The percentage of crosslinking efficiency in panel H was calculated as mean+SD of 3 independent experiments. The highest increase in crosslink was observed when both R1008I and W483R were present, about 70% of wild type. (D) Scc2 D369C and *smc3QQ* K1004C were crosslinked *in vivo* in the presence of R1008I and W483R or both using BMOE. The first lane shows the crosslink of wild type Smc3. Immunoprecipitated on Scc1-PK, separated on gradient gel and analysed by western blot. (E) The percentage of crosslinking efficiency in panel D was calculated as mean+SD of 3 independent experiments. Smc3 R1008I increased the crosslink of smc3QQ with Scc2 to about 70% of wild type, regardless of Smc3 K1004C. (F) Scc2 E822K L937F (red circles) promotes cell proliferation of smc3QQ R1008I mutant (green circles), nearly to wild type (red rectangles). (G) Calibrated ChIP-seq profiles of *Smc3, smc3QQ, smc3QQ R1008I*, in the presence of *WT Scc2* or *Scc2 E822K L937F*. (D) Recombinant tetramer of cohesin, Smc3 or smc3QQ, Smc1, Scc1 and Scc3 was incubated with Scc2 or Scc2 E822K L937F. ATP was added to initiate the reaction and the reaction rate was measured as the change in absorption at 360nm over time. (E) ATPase analysis of recombinant tetramers of cohesin, Smc3 or smc3QQ R1008I, Smc1, Scc1 and Scc3 was incubated with Scc2 or Scc2 E822K L937F.

We, therefore, expected that the defective Scc2-dependent ATP activity of Smc3QQ can also be improved by *R1008I W483R*. To our surprise, *W483R R1008I* mildly increased the Scc2- dependent ATPase activity of Smc3QQ when DNA is absent (Fig 6E), although it restored the DNA association of Smc3QQ to a wild-type level (Fig 1F). It revealed that the main role of these mutations is to enhance the interaction of Smc3QQ with Scc2 and form the Scc2/pre-E-cohesin complex. This complex can be recruited to DNA, further enhancing ATP hydrolysis. This explains why *W483R R1008I* can fully recover the loading of Smc3QQ with mildly increased Scc2-dependent ATPase activity in the absence of DNA.

### Smc3 acetylation might impair cohesin’s intrinsic ATPase activity

The above experiments revealed that the Scc2-dependent ATPase activity of Smc3QQ is not much increased by the suppressor mutation *W483R R1008I* although the Scc2/Smc3QQ interaction is greatly improved. This raised a possibility that the intrinsic ATPase activity might be affected by *smc3QQ* or Smc3 acetylation. Consistently, we also identified 16 mutations on the Smc3 head domain, including Q-loop, signature motif, and ATPase binding pocket, which suppress the growth defect of *smc3QQ R1008I* (Fig S6F). Three of these mutations (*K158E, Q1143R*, and *C1183S*) are sufficient to rescue the lethality of *smc3QQ* on their own, although the growth of these cells was poor. The additional mutation *R1008I* greatly improved their growth. The remaining mutations cannot support the growth of *smc3QQ* cells by themself, but significantly alleviate the growth defect of *smc3QQ R1008I* cells. The distributions of these head mutations in the regions responsible for ATP binding/hydrolysis support our hypothesis that Smc3 acetylation might also impair the intrinsic ATPase activity. The activity requires ATP binding/head engagement and ATP hydrolysis. Since ATP binding-mediated head engagement is not impaired by *smc3QQ* (Fig 2C), thus, we proposed that enhancing ATP hydrolysis would remedy the growth defect of *smc3QQ*. To test this, we examined the effect of *scc2 E822K L937F (scc2EKLF)*, a hypermorphic Scc2 allele ^4^, on the growth of *smc3QQ* or *smc3QQ R1008I* mutant. Strikingly, although *scc2EKLF* cannot suppress the lethality of *smc3QQ* (data not shown), it greatly promotes cell proliferation of *smc3QQ R1008I* mutant (Fig 6F). Further tetrad dissection revealed that either *scc2 E822K* or *L937F* is sufficient to improve the growth of *smc3QQ-R1008I* cells (Fig S6G). To evaluate the effect of *scc2EKLF* on the DNA association of Smc3QQ and Smc3QQ R1008I, we ectopically expressed Smc3, Smc3QQ, or Smc3QQ-R1008I in wild-type *SCC2* or *scc2EKLF* cells (Fig 6G). Consistent with our previous results, calibrated ChIP-seq revealed serious defects in the DNA association of Smc3QQ and Smc3QQ R1008I. Although *scc2EKLF* barely affected the loading of Smc3QQ, it significantly elevated the DNA association of Smc3QQ R1008I from centromere to chromosomal arms, about ∼40% to ∼90% of wild-type Smc3. Intriguingly, the ATP hydrolysis of Smc3QQ and Smc3QQ R1008I were dramatically increased by *scc2EKLF* (Fig 6H and 6I). Together with the distribution of suppressor mutations on the head subdomains involving ATP hydrolysis, it implied that Smc3 acetylation impairs its intrinsic ATPase activity and the hypermorphic Scc2 allele can compensate for this defect. However, the restoration of ATP hydrolysis of Smc3QQ is insufficient for its loading and still requires *smc3 R1008I*, although *R1008I* barely contributes to the ATPase activity of Smc3 or Smc3QQ (Fig S6H). Considering the role of *smc3 R1008I* in the interaction of Scc2 with Smc3QQ CC, we inferred that this mutation would facilitate the interaction of Scc2EKLF with Smc3QQ CC and form the DNA-clamp compartment. This explains why R1008I is still required for Scc2EFLK to suppress the loading defect of *smc3QQ*.

### ATP-dependent head engagement promotes DNA clamping

We next surveyed how ATP binding-dependent head engagement regulated the conformational dynamics of the Scc2/cohesin loading complex. To this end, we examined whether the four Scc2/cohesin interfaces are affected by ATP hydrolysis mutants s*mc3 E1155Q* and s*mc1 E1158Q*, which blocks ATP hydrolysis and is likely to preserve the cohesin configuration with engaged heads. We found that the ATP hydrolysis mutants barely altered the crosslinks of Smc1 E1102C/Scc2 T181C and Smc3 Q67C/Scc2 T1175C (Fig 7A, 7B, and S7A). These results are consistent with the Scc2/J-cohesin or pre-E-cohesin configurations, in which the heads are disengaged. The *smc3 E1155Q* doubled the Smc3 S72C/Scc2 E819C crosslink and greatly improved the Smc3 K1004C/Scc2 D369C crosslink by about 10 times. This indicates that the interface of Smc3 CC with Scc2 is driven by ATP binding and head engagement. It suggested that the head engagement would orientate Smc3 CC and facilitate its interaction with Scc2. Because this interface is critical for the DNA-clamp compartment (Shi et al. 2020; Higashi et al. 2020; Collier et al. 2020), we assumed that head engagement has an important role in regulating this compartment and subsequent DNA entrapment. To examine this compartment *in vivo*, two cysteine substitutions *E819C* and *D369C* were introduced into Scc2 and the corresponding *S72C* and *K1004C* were also introduced Smc3. To investigate the effect of head engagement on the DNA-clamp compartment, we examine the crosslinks in the presence of *smc3 E1155Q*. As expected, the crosslink of Smc3 K1004C/Scc2 D369C is inefficient with wild-type Smc3 and dramatically increased by *smc3 E1155Q* to a similar level of the Smc3 S72C/Scc2 E819C crosslink (Fig 7C). Strikingly, an extra crosslink band appeared at the top of Western blot in the *smc3 E1155Q* strain bearing both cysteine pairs. This band, produced by double crosslinking, represents the DNA-clamp compartment, as seen in the Scc2/E-cohesin cryo-EM structure. The double crosslink product is barely detectable in *SMC3* cells (Fig 7C). All these data uncover one of the essential roles of ATP-dependent head engagement, which modulates the orientation of the Smc3 CC and promotes its interaction with Scc2. This is essential for closing the DNA-clamp compartment and permitting clamping DNA.

**Fig 7:**
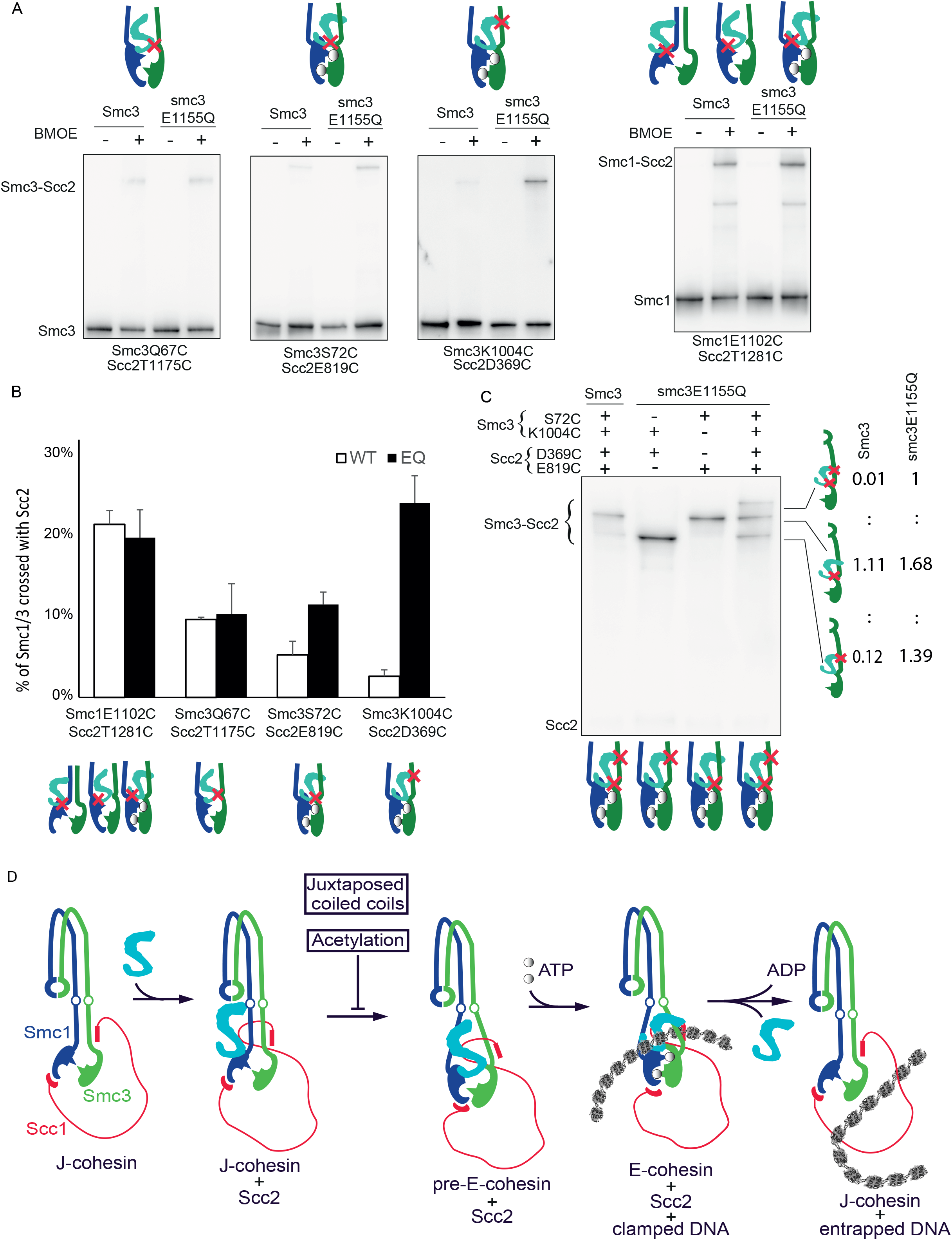
ATP binding facilitates formation of the DNA-clamp compartment. (A) Effect of Smc3 ATP hydrolysis mutant (*smc3* E1155Q) on the crosslinks at different Scc2/cohesin interfaces. Wild type or E1155Q mutant versions of Smc3 S72C, Smc3 K1004C, Smc3 Q67C (right panel) or Smc1 E1102C/Scc2 T1281C in cells with ectopically expressed *Smc3*-PK or *smc3 E1155Q*-PK (left panel) were *in vivo* BMOE crosslinked to their respective cysteine substituted Scc2. Immunoprecipitated on Scc1-PK (right panel) or Smc3-PK (left panel), separated on gradient, gel and analysed by western blot. Crosslinking efficiency was quantified using band intensity on western blot. (B) The percentage of crosslinking efficiency in panel A was calculated as mean+SD of 3 independent experiments. *smc3 E1155Q* increased the crosslinking efficiency of Scc2 with Smc3 S72C and Smc3 K1004C. The highest increase of about 10-fold was seen in the crosslink Smc3 K1004C/Scc2 D369C. (C) DNA clamping compartment formed by the interfaces of Smc3 S72/Scc2 E819 and Smc3 K1004/Scc2 D369. Simultaneous *in vivo* BMOE crosslink of Smc3 S72C/Scc2 E819C and Smc3 K1004C/Scc2 D369C with *WT Smc3* or *smc3 E1155Q* was performed. Immunoprecipitation was done using Scc1-PK, separated on gradient gel, and Scc2 was analysed by western blot. The single and double crosslink products were quantified using band intensity on western blot. Their ratios were indicated on the left. (D) Conformational dynamics of cohesin Scc2 loading complex. Scc2 interacts in J-cohesin with Smc1’s head. Then the interaction of Scc2/Smc3 heads induces the unzipping of cohesin CC and orientates heads to facilitate the formation of ATP binding pocket (pre-E-cohesin). ATP binding leads to head engagement (E-cohesin) and promotes the assembly of the DNA clamping compartment. DNA clamping triggers ATP hydrolysis and the opening of one of the cohesin interfaces, allowing DNA to be entrapped in cohesin’s J-K compartment. The transition from the configuration of Scc2/J-cohesin to that of Scc2/pre-E-cohesin is inhibited by Smc3 acetylation and juxtaposed CCs.

## Discussion

### Identification of a novel Scc2/pre-cohesin complex

Scc2/4 complex is a cohesin loader, essential for its various functions, such as sister chromatid cohesion and loop extrusion ^35–37^. During loading, Scc2 is believed to form a complex with cohesin and activates its ATPase (Murayama & Uhlmann 2014; Petela et al. 2018). Cohesin also has two configurations: J-cohesin and E-cohesin. J-cohesin is responsible for the topological entrapment of DNA in the J-K compartment, which is the final step of loading ^12^. However, the juxtaposed CCs in J-cohesin impose a restraint on head engagement, therefore, this configuration is incompatible with ATP binding and hydrolysis. E-cohesin is known as a crucial configuration for ATP hydrolysis and the loading reaction (Arumugam et al. 2003; Hu et al. 2011). Recently, several *in vitro* Cryo-EM demonstrated that Scc2 can bind both cohesin complexes^11,13–15^. Interestingly, the Scc2/E-cohesin complex can clamp DNA in a small compartment encircled by Scc2 and Smc3 CC. This structure would reflect an essential step of the loading reaction. Therefore, it is likely that Scc2 can interact with J-cohesin, load it to DNA, and form the Scc2/E- cohesin DNA clamp configuration (Fig S1A). The subsequent ATP hydrolysis drives DNA to be entrapped in the J-K compartment associated with kleisin and the heads of J-cohesin. However, it is unclear how Scc2 loads the J-cohesin onto DNA to form the DNA-clamped Scc2/E-cohesin complex.

In this study, we confirmed the existence of both configurations *in vivo*. Interestingly, we identify another Scc2/cohesin configuration (Fig 3H), which is defined by two interfaces between Scc2 and cohesin, Scc2 T1281/Smc1 E1102 and Scc2 T1175/Smc3 Q67. Several lines of evidence derived from this study demonstrated that this configuration is distinguishable from the other two. First, this configuration is incompatible with J-cohesin because juxtaposed CCs impose a structural restraint on the contact of Scc2 T1175/Smc3 Q67. This is confirmed by sequential crosslinks (Fig 4C). Using the same strategy, we demonstrated that head engagement also prevents the formation of this interface (Fig 4D). These results are consistent with the prediction based on our modelled structure: Scc2 interacts cohesin with separated CCs and disengaged heads. Therefore, we proposed that the interaction of Scc2 T1175/Smc3 Q67 represents a novel configuration, Scc2/pre-E-cohesin. This hypothesis is further supported by the distinctive sensitivities of these three configurations to the acetylation mimicking mutation *smc3QQ* and ATP hydrolysis mutation *smc3 E1155Q*. The Scc2/pre-E-cohesin and Scc2/E- cohesin, but not Scc2/J-cohesin, are seriously defective with *smc3QQ* (Fig 5B, 5G, 5I, and 5K). It suggested that the interaction of Scc2 with J-cohesin is the first step in loading. Also, enhancement of only the Scc2/E-cohesin by *smc3 E1155Q* (Fig 6B) suggests the occurrence of Scc2/J-cohesin and Scc2/pre-E-cohesin before the Scc2/E-cohesin. Together, these results support the notion that Scc2/pre-E-cohesin is a novel configuration and acts as a transitional step during loading (Fig 7D).

### The Scc2/pre-cohesin configuration promotes the transition from Scc2/J-cohesin to DNA-clamped Scc2/E-cohesin during loading

Our observation that disruption of the Scc2/pre-cohesin complex by Smc3 acetylation-mimicking mutant *smc3QQ* prevents its DNA association demonstrates an essential of this configuration in loading. How does the Scc2/pre-cohesin configuration promote the loading reaction? As suggested by a Cyro-EM study, Scc2 interacts with J-cohesin^15^, which initiates the loading reaction. However, the misaligned heads in this Scc2/J-cohesin configuration are incompatible with ATP binding and the juxtaposed CCs prevent the formation of the DNA-clamped compartment. In the Scc2/pre-cohesin configuration, the interaction of Scc2 with both heads brings them in close proximity, facilitating ATP binding and head engagement. Moreover, the association of Scc2 with both heads also drives the unzipping of cohesin CCs, permitting the interaction of Scc2 with Smc3 CC and the formation of the compartment for the DNA clamp. On the other hand, Scc2 can also interact with E-cohesin, and DNA is clamped in the Scc2/Smc3 compartment^11,13,14^. However, if this configuration forms prior to cohesin/DNA association, Scc2 can trigger ATP hydrolysis before loading cohesin onto DNA, which will cause the disassembly of this configuration. Another problem is that the engaged heads promote the interaction of Scc2 with Smc3 heads and CC (Fig 7A), which encloses the DNA-clamp compartment and prevents DNA entry into this compartment. Unlike the Scc2/E-cohesin configuration, the heads in the Scc2/pre-E-cohesin complex are disengaged, which is unable to bind ATP. This structural feature prevents premature ATP hydrolysis before cohesin is loaded onto DNA. Importantly, although Scc2 interacts with the Smc3 head in the Scc2/pre-E-cohesin configuration, the unengaged heads inhibit the interaction between Scc2 and Smc3 CC and keep this interface open. This creates an entry gate for DNA to enter the DNA-clamp compartment. The subsequent DNA binding drives Scc2/Smc3 CC interaction and the close of this gate. These structural features of the Scc2/pre-E-cohesin configuration permit Scc2 to load J-cohesin and form the DNA-clamped Scc2/E-cohesin configuration.

### Regulation of the loading reaction by Smc3 acetylation

Acetylation of Smc3 K112 K113 is critical to prevent DNA disassociation of cohesin by antagonising Wapl’s releasing activity ^18,22,24,39^. Ironically, this modification also precludes Scc2- mediated DNA association as revealed by studying its mimicking form *smc3QQ* ^10,29^. These observations suggest that Smc3 K112 K113 plays essential roles in both loading and release. How do these two residues function in opposite processes? One explanation is that the opening of the Smc3/Scc1 interface promoted by K112 K113 might produce a gate to allow DNA entry in loading and exit during release. Although the acetylation-mimicking mutant, *smc3QQ*, has been widely used to study Smc3 acetylation, there is no evidence to demonstrate that K112Q K113Q can maintain cohesion as acetylated Smc3 K112 K113. This is because Smc3QQ cannot be loaded onto DNA in the first place. The identified mutation in this study, *smc3 W483R R1008I*, suppresses the lethality of *smc3QQ*, proving that *smc3QQ* can functionally copy Smc3 acetylation to prevent releasing activity and maintain cohesion in the presence of Wapl. Moreover, this mutation largely restores the DNA association of Smc3QQ but has little effect on release, suggesting that Smc3 K112 K113 activate loading and release through distinctive pathways.

How do Smc3 K112 K113 promote loading? The Cryo-EM analysis of Scc2/E-cohesin/DNA configuration revealed that these two residues are in close proximity to DNA or two conserved negatively charged residues E821 E822 of Scc2 ^11,13^. Therefore, two hypotheses based on the charge of Smc3 K112 K113 were proposed: positively charged K112 K113 acts as a DNA sensor by interacting with negatively charged DNA or promotes the interaction with Scc2 by forming electrostatic forces with Scc2 E821 E822. Thus, neutralisation of their charge by acetylation would weaken these interactions and inhibit loading. Although we cannot exclude these possibilities, several lines of evidence suggest that these interactions might not account for the essential role of K112 K113 in loading. First, DNA can significantly enhance the Scc2- dependent ATPase activity of Smc3QQ (Fig 2B), suggesting that neutralising the positive charges of Smc3 K112 113 does not seriously impair cohesin/DNA interaction. On the other hand, if electrostatic forces, created by Smc3 K112 K113 and Scc2 E821 E822, are essential for Smc3/Scc2 interaction and loading, it is expected that neutralising charges of Scc2 E821 E822 should also cause loading defects. However, unlike *smc3QQ, scc2 E821Q/E822Q* barely affects cell growth and sister chromatid cohesion (data not shown). In contrast, reversing the positive charge of Scc2 E822 leads to a gain-of-function allele of Scc2 (*scc2-E822K*), which can efficiently stimulate the ATPase activity of Smc3QQ and Smc3. Interestingly, this allele, together with *R1008I*, suppresses the loading defect and lethality of Smc3QQ (Fig S6G). These results indicated that DNA or Scc2 is still able to interact with cohesin in the loading process even though the positive charge of K112 K113 is neutralised. We, therefore, concluded that the interaction of Smc3 K112 K113 with DNA or Scc2 E821 E822 might not be a major contributor to loading.

This study revealed that *smc3QQ* shows a minimal effect on Scc2/Smc1 interface, so *smc3QQ* is still able to form a complex with Scc2 interaction, which is expected in the Scc2/J-cohesin configuration. This is not surprising since there is no direct contact between Scc2 and Smc3 in this configuration. However, we found that *smc3QQ* dramatically decreases the interaction between Scc2 and Smc3 Q67. It suggests that the acetylation mimicking mutant impairs the Scc2/pre-E-cohesin configuration and prevents the conformational transition from Scc2/J- cohesin to Scc2/E-cohesin. Consistently, the mutation *smc3 W483R R1008I* which improves the interaction of Scc2 with Smc3QQ also greatly alleviates the loading defect of Smc3QQ. This suggests that the main role of Smc3 K112 K113 in loading is to facilitate the formation of Scc2/pre-E-cohesin configuration. However, how these residues influence this configuration still requires further investigation.

To our surprise, although the interaction of Scc2 with Smc3QQ and the loading of Smc3QQ are greatly improved by s*mc3 W483R R1008I*, this suppressor mutation has a mild effect on the reduced DNA-independent ATPase activity of Smc3QQ. This suggested that Smc3 acetylation might also impair the intrinsic ATPase activity. In agreement with this hypothesis, many mutations suppressing the growth defect of *smc3QQ* were found in the regions involving ATP binding and hydrolysis. These mutations might improve the intrinsic ATPase activity of *smc3QQ*.

Moreover, we found a hypermorphic allele of Scc2, together with *smc3 R1008I*, can efficiently suppress the loading defect of *smc3QQ*. Consistently, a mutation in the Smc1 head greatly improves the intrinsic ATPase activity and suppresses the lethality of *smc3 K113Q* (Koshland, personal communication). These results suggested that Smc3 K112 K113 would also contribute to the intrinsic ATPase activity.

### Juxtaposed Smc CCs block the Scc2/pre-E-cohesin and Scc2/E-cohesin configurations

In this study, we identified numerous mutations across the Smc3 CC suppressing the lethality of *smc3QQ* and remedying its loading defects. This indicates that cohesin CCs do not only act as structural blocks but also actively regulate cohesin functions, such as loading in this case. Recent evidence revealed that SMC CCs are juxtaposed with each other through extensive interactions ^40–42^. This configuration will pose a structural barrier for Smc3/Scc2 interaction and ATP binding. We demonstrated that the chemical stabilisation of juxtaposed CCs prevents Scc2/Smc3 Q67 interaction. Thus, the juxtaposed CCs must be unzipped, at least from “necks” to “heads”, for assembly of the Scc2/pre-E-cohesin complex and further ATP binding/hydrolysis. Also, Cryo-EM revealed that Scc2 and the Smc3 head-end CC create a compartment for DNA clamping, which also requires the unzipping of juxtaposed CCs. Consistently, two main suppressor mutations *E199A* and *R1008I* localise to the end of Smc3 CC close to its ATPase head, which might weaken the interaction between cohesin CCs. We infer that this weakening would facilitate the unzipping of juxtaposed coiled coils. Interestingly, this effect can be further enhanced by additional mutations occurring on the Smc3 CC. The coiled coils of SMC proteins are not made of a continuous α-helix and are broken into several fragments of about 50 residues by conserved disruption ^43^. Remarkably, most CC suppressor mutations are found to locate at joints of α-helix fragments, which might contribute to unzipping juxtaposed CCs by altering the orientation of the α-helix. Our investigation revealed that *W483R R1008I* greatly improved the interaction of Scc2 with Smc3QQ Q67. This improvement permits the assembly of the Scc2-pre- E-cohesin complex and enables the loading. All these data suggest that the juxtaposition of cohesin CCs poses a structural barrier for Scc2/Smc3 head interaction and prevents Scc2/pre-E- cohesin configuration, which hinders loading.

### ATP-dependent head engagement regulates the DNA clamp

During loading, DNA is revealed to be clamped in the small compartment enclosed by two interfaces: Scc2/Smc3 head and Scc2/Smc3 CC. The questions are how DNA enters this compartment and how ATP regulates this process. This study revealed that the Scc2/Smc3 CC interaction relies on ATP-dependent head engagement (Fig 7B). This suggested that ATP binding-triggered head engagement orientates Smc3 CC to permit its interaction with Scc2. However, the interaction of Scc2 with the Smc3 head would not require head engagement, for example, in the Scc2/pre-E-configuration (Fig 7B). This suggested that the Scc2/Smc3 CC will be the entry gate for DNA and ATP promotes the close of this gate, allowing DNA to be entrapped in this compartment. Since the configuration of the Scc2/cohesin complex is highly dynamic, understanding how this conformational dynamic is regulated during loading and loop extrusion will provide mechanistic insights into both processes.

## Methods

### Yeast strains and growth conditions

All yeast strains were of W303 background (Table S2). Standard yeast genetic techniques and transformation were used to construct these strains. Integration of target genes into the *MET15* locus using the CRISPR system as previously described^44^.

Unless otherwise stated, yeast cells were grown in the rich medium YEP + 2% glucose (YEPD) at 25°C with a rotation of 2000 rpm (New Brunswic Innova 43R, Eppendorf). To arrest cells in late G1 by overexpressing non-degradable Sic1(9m) ^22^, exponentially grown *bar1Δ* cells in YEP + 2% raffinose (YEPR) were treated with 0.1 µg/ml α-factor. One hour before release, galactose of a final concentration of 2% was added to induce the expression of Sic1(9m). to release cells from α-factor arrest, cells were harvested by centrifuge (3500rpm × 2 minutes) and washed with YEPD twice. Cells were resuspended into YEPD containing 0.1 mg/ml of pronase and agitated for 60 min at 25°C.

Yeast tetrad dissection was performed using a Singer dissection microscope MSM 400.

## Method Details

### Screening for suppressors of *scc2-45 wpl1Δ*

Forty independent colonies of the parental strain (*scc2-45 wpl1Δ*) were cultured in 5ml YEPD overnight at 25°C. Cells collected from 1ml culture were spread ontothree3 plates and incubated at 32°C until colonies appeared. A colony was picked from each and confirmed robust growth at 32°C by streaking on YEPD plate. Eighty-eight revertants of Scc2 were identified by sequencing the Scc2 gene amplified from genomic DNA using PCR. Genetic linkage of the rest mutations with known cohesin submits (*SMC1, SMC3, SCC1, SCC3, SCC4, PDS5* and *ECO1*) were examined by tetrad dissection. Twenty-seven were tightly linked to SMC3 and DNA sequencing revealed numerous substitutions of just two amino acids, namely E199 and R1008.

### Screening for suppressors of *smc3QQ R1008I*

Random mutations were introduced into the *smc3 R1008I* gene using two-round error-prone PCR. In the first round of PCR, MnCl_2_ was added to a final 30 μM to induce A/T to G/C transition. In the second round of PCR, 1 μl of the first-round PCR product was used as the template DNA, and dITP was added to a final 30μM to induce G/C to A/T transition. Both PCRs were carried out by standard PCR using G2 DNA Polymerase (Promega). The mutated smc3 R1008I and linearized YCplac111 bearing Smc3 promoter or terminator plus 50bp sequences homologous to Smc3 ORF at each end were co-transformed into smc3-42 cells. The transformants containing suppressor mutations were selected on SD -Leu plates at 37°C. The YCplac111 plasmids were recovered, and the mutations were identified by DNA sequencing. Sixteen suppressor mutations on the Smc3 head and twenty-four on Smc3 CC were verified by replacing endogenous Smc3 with the reconstituted mutations.

### *In vivo* BPA crosslink

*In vivo* BPA crosslink was performed as previously described ^45^. Yeast cells were transformed with pBH61 expressing Ec TyrRS and Ec tRNACUA, and YEplac181 containing Smc1 or Smc3 gene with a TAG mutation. These cells were grown in SD -Leu -Trp media containing 1mM BPA at 30°C until the OD reached 0.6 per mL. The BPA stocking solution (1M) was made by dissolving BPA in fresh 1N NaOH solution. The cells (40 OD) were collected, resuspended in 1mL ice cold 1× PBS and transferred into an 8-well plate. The plate was placed on a bed of ice and put into a UV crosslinker (Spectrolinker XL-1500W crosslinker, Spectronics Corporation) equipped with a 352nm wavelength emitting bulb (Sankyo Denki Co. Ltd). The cells were subjected to UV light at 352nm for 3× 1 minute with a 5-minute rest interval.

### *In vivo* BMOE crosslink

A stock solution of 125mM BMOE and dissolved at 37°C with shaking. First, yeast cells were arrested in later G1. Then 30 OD of cells were collected and washed with 500 µl of PBS twice before re-suspending in 1 ml of PBS. 25µL of 125mM BMOE solution dissolved in DMSO was added to the cell suspension and incubated on ice for 6 minutes. The crosslinking reaction was stopped by adding 3µl of 1M DTT. The cells were then washed with 1mL of 5mM DTT in PBS and used for protein extraction.

### Co-immunoprecipitation

Proteins were extracted from 30 OD of cells in 500 µl of lysis buffer (0.05M Tris pH 7.5, 0.15M NaCl, 5mM EDTA pH 8.0, 0.5% NP-40, 1mM DTT, 1mM PMSF, and cOmplete™ Protease Inhibitor Cocktail tablet). The crude lysis was added with 3 µl of 1mg/ml antibody and rotated at 4°C for 1 hour. Then 30 µl of Dynabeads (Thermo Fisher) was added and rotated at 4°C overnight. The Dynabeads were washed three times using 1 ml of lysis buffer per wash with a 5- minute incubation with rolling at 4°C between each wash. The lysis buffer was removed and the IPed proteins were released in 50µL of 1× LDS sample buffer (NuPAGE^®^ Life Technologies) by heating at 65°C for 5 minutes.

### TEV proteinase cleavage

TEV cleavage was performed after the wash step during the immunoprecipitation. The Dynabeads were then washed once with 500μL of digestion buffer (25mM Tris pH 8.0, 150mM NaCl, and 2mM 2-Mercaptoethanol). The beads were then incubated for 16 hours at 20°C with shaking in 30 μL of digestion buffer supplemented with 10 μl of TEV protease (Sigma). The samples were mixed with 10μl of 1× LDS sample buffer and heated at 65°C for 5 minutes.

### Protein gel electrophoresis and Western blotting

The crosslinked samples in 1X LDS sample buffer were separated using 3-8% Tris-acetate gels (NuPAGE^®^ Life Technologies). For single crosslinks, SDS-PAG lasted 150 minutes with 150 V. For double crosslinks, SDS-PAG lasted 210 minutes with 150 V. The proteins were then transferred onto 0.2μm nitrocellulose using the Mini Trans-blot^®^ Cell system (Bio-Rad).

| <u>Primary antibody</u> | <u>Immunogen</u> | <u>Origin</u> | <u>Dilution</u> |
| --- | --- | --- | --- |
| V5 (Bio-Rad) | PK | Mouse | 1:3,000 |
| 3F10 (Roche) | HA | Rat | 1:3,000 |
| 4A6 (Millipore) | MYC | Rat | 1:3,000 |
| M2 (Sigma) | FLAG | Rat | 1:3,000 |
| PGK1 | PGK1 | Mouse | 1:5,000 |

#### Secondary antibody

Goat anti-Mouse IgG (H/L):HRP (Bio-Rad) (1:3,000)

To visualize the proteins, the luminant signal was developed using Immobilon^™^ Western Chemiluminescent HRP substrate (Millipore) and detected using G:Box Chemi-XX9 (GENESYS). The density of each band was measured using GeneTools (GENESYS).

### Calibrated ChIP-sequencing

Calibrated ChIP-sequencing was performed as previously described ^29^. Forty-five mL of 15 OD_600_ units of exponentially grown cells and 5 OD_600_ units of reference cells *C. glabrata* were mixed with 4mL of fixative (50mM Tris-HCl, pH 8.0; 100mM NaCl; 0.5mM EGTA; 1mM EDTA; 30% (v/v) formaldehyde) and rotated at RT for 30min. Then 2mL of 2.5M glycine was added and incubated at RT for 5min. The fixed cells were collected by centrifugation at 3,500rpm for 2min and washed with 50ml of ice-cold 1X PBS. The cells were then resuspended in 300μL of ChIP lysis buffer (50mM Hepes-KOH, pH 8.0; 140mM NaCl; 1mM EDTA; 1% (v/v) Triton X-100; 0.1% (w/v) sodium deoxycholate; 1mM PMSF; 1 tablet/25mL protease inhibitor cocktail (Roche)) and an equal amount of acid-washed glass beads (425-600μm Sigma) added. Cells were disrupted using a FastPrep^®^-24 benchtop homogeniser (M.P. Biomedicals) at 4°C (3x 60s at 6.5m/s or until >90% of the cells were lysed as confirmed by microscopy).

The isolated soluble fraction was subjected to a sonication using a bioruptor (Diagenode) for 30min in bursts of 30s on and 30s off at a high level in a 4°C water bath. Under this condition,. Cell debris was removed by centrifugation at 13,200rpm at 4°C for 20min and the supernatant containing sheared chromatin with a size range of 200-1,000bp was mixed with 700μL of ChIP lysis buffer. The samples were pre-cleared with 30μL of protein G Dynabeads (Invitrogen) for 1h at 4°C. 80μL of WCE was taken. The remaining supernatant was mixed with 5μg of anti-PK antibody Bio-Rad) and rotated overnight at 4°C. 50μL of protein G Dynabeads were added and rotated at 4°C for 2h. The beads were washed 2x with ChIP lysis buffer, 3x with high salt ChIP lysis buffer (50mM Hepes-KOH, pH 8.0; 500mM NaCl; 1mM EDTA; 1% (v/v) Triton X-100; 0.1% (w/v) sodium deoxycholate; 1mM PMSF), 2x with ChIP wash buffer (10mM Tris-HCl, pH 8.0; 0.25M LiCl; 0.5% NP-40; 0.5% sodium deoxycholate; 1mM EDTA; 1mM PMSF) and 1x with TE pH 7.5. The immunoprecipitated chromatin was then released in 120μL of TES buffer (50mM Tris-HCl, pH 8.0; 10mM EDTA; 1% SDS) by heating at 65°C for 15min and the collected supernatant termed the IP sample. The WCE extracts were mixed with 40μL of TES3 buffer (50mM Tris-HCl, pH 8.0; 10mM EDTA; 3% SDS). The crosslink of WCE and IP samples were reverted at 65°C overnight. RNA was digested with 2μL RNase A (10mg/mL; Roche) for 1h at 37°C and protein was degraded with 10μL of proteinase K (18mg/mL; Roche) for 2h at 65°C. DNA was purified using ChIP DNA Clean and Concentrator kit (Zymo Research).

### Preparation of sequencing libraries

Sequencing libraries were prepared using NEBNext^®^ Fast DNA Library Prep Set for Ion Torrent^™^ Kit (New England Biolabs) following the manufacturer’s instructions. Briefly, 10- 100ng of purified DNA was subjected to end repair and ligated with the Ion Xpress^™^ Barcode Adaptors. Fragments of 300bp were then selected using E-Gel^®^ SizeSelect^™^ 2% Agarose gels (Life Technologies) and amplified with 6-8 PCR cycles. The DNA concentration was then determined by qPCR using Ion Torrent DNA standards (Kapa Biosystems) as a reference. 12-16 libraries with different barcodes then pooled together to a final concentration of 350pM and loaded onto the Ion PI^™^ V3 Chip (Life Technologies) using the Ion Chef^™^ (Life Technologies). Sequencing was then completed on the Ion Torrent Proton (Life Technologies), typically producing 6-10 million reads per library with an average read length of 190bp.

### Data analysis, alignment and production of BigWigs

Data analysis was performed on the Galaxy platform ^46^. The quality of the reads was assessed using FastQC (Galaxy tool version 1.,0.0) and the low-quality bases were removed as required using ‘trim sequences’ (Galaxy tool version 1.0.0). Generally, this involved removing the first 10 bases and any bases after the 200^th^ but trimming more or fewer bases may be required to ensure the removal of kmers and that the per-base sequence content is equal across the reads. Reads shorter than 50 bp were removed using Filter FASTQ (Galaxy tool version 1.0.0, minimum size: 50, maximum size: 0, minimum quality: 0, maximum quality: 0, maximum number of bases allowed outside of quality range: 0, paired end data: false) and the remaining reads aligned to the necessary genome(s) using Bowtie2 (Galaxy tool version 0.2) with the default (--sensitive) parameters (mate paired: single-end, write unaligned reads to a separate file: true, reference genome: SacCer3 or CanGla, specify read group: false, parameter settings: full parameter list, type of alignment: end to end, preset option: sensitive, disallow gaps within *n-* positions of read: 4, trim *n*-bases from 5’ of each read: 0, number of reads to be aligned: 0, strand directions: both, log mapping time: false).

To generate alignments of reads that uniquely align to the *S. cerevisiae* genome, the reads were first aligned to the *C. glabrata* (CBS138, genolevures) genome and retrieved the unaligned reads. These unaligned reads were then aligned to the *S. cerevisiae* (sacCer3, SGD) genom,e and the resulting BAM file was converted to BigWig as (Galaxy tool version 0.1.0) and the reads are normalised as counts per million (CPM). Similarly this process was done with the order of genomes reversed to produce alignments of reads that uniquely align to *C. glabrata*.

### Visualisation of ChIP-seq profiles

The BigWigs were visualised using the IGB browser ^47^. To calibrate the ChIP profile, the track was multiplied by the samples occupancy ratio (OR) using the graph multiply function.

The Bigwigs file was used to calculate the average occupancy of the 10kb region around all 16 centromeres through the tool “computeMatrix” and OR was used as the scale factor. A BED file named “yeast CDEIII” is used to define the centres of all the 16 CDEIII in this calculation.

The accession number for the calibrated ChIP-seq data (raw and analysed) reported in this paper is GEO: GSE217833.

### Protein purification of the cohesin and Scc2 complexes

The codons-optimized Smc1/8xHis-Smc3 or smc3QQ/Scc1 2xStrepII/Scc3 and GFP- ΔΝ132-*scc2*-1xStrepII were into bacmids and transfected into Sf9 cells using Fugene HD reagent (Promega). The generated viruses were infected into Sf9 cells, and the cells were cultured at 27 °C for 72h in Insect-XPRESS protein-free medium with L-glutamate (Lonza). To express the proteins, 500ml of SF-9 insect cells with a cell density of ∼3 million/ml were infected with the appropriate baculovirus stock in a 1/100 dilution and harvested when lethality (assayed by the trypan blue test) reached no more than 70-80%. Cell pellets were quickly frozen in liquid nitrogen and stored at -80°C. The thawed pellets were suspended in ∼65-70ml of HNTG lysis buffer (25mM Hepes pH 8.0, NaCl 150mM, TCEP-HCl 1mM and Glycerol 10%. The suspension was immediately supplemented with 2 dissolved tablets of Roche Complete Protease (EDTA-free), 75µg of RNAse I and 7µl of DNAseI (Roche, of 10U/µl stock). The cells were then sonicated at 80% amplitude for 5s/burst/35ml of suspension using a Sonics Vibra-Cell (3mm microtip). In total 5 bursts were given for every 35ml half of the 70ml suspension (the sonication was always performed in ethanolised ice). Cell debris were removed using Ti45 fixed angle rotor (45,000rpm x 45 mins). A final concentration of 2mM EDTA was added to the cleared extract and loaded to a 2×5ml StrepTrap HP (Fisher Scientific) column at 1ml/min in an ÄKTA Purifier 100. Washed with HNTG containing 1mM PMSF and 2mM EDTA (HNTGPE) with a flow speed of 1ml/min until ΔΑU_280nm_ is reduced to ∼0. The protein was eluted using HNTGPE+20mM desthiobiotin (Fisher Scientific) at 1ml/min. Peak fractions analysed using SDS-PAGE were pooled and were further purified in a Superose 6 Increase 10/300 (VWR) using HNTG as a running buffer (free of EDTA/PMSF). The resulting peaks were again analysed using SDS-PAGE and the concentration was determined by measuring A280 using Nanodrop. Protein was aliquoted and stocked typically in concentrations ranging from 1 to 3mg/ml.

### ATPase assay

ATPase activity was determined as described in the protocol of the EnzChek phosphate assay kit (Invitrogen). The tetramer cohesin of a final concentration of 50nM was dissolved with 50mM NaCl. If dsDNA is presented, 700nM 40bp dsDNA was used. The reaction was initialised by adding ATP to a final concentration of 1.3mM (final reaction volume: 150ul). The OD at 360nm was monitored every 30s for 90min using a PHERAstar FS. ΔΑU at 360nm was translated to Pi release using an equation derived by a standard curve of KH_2_PO_4_ (EnzChek kit). Rates were calculated from the slope of the linear phase (first 10min). At least two independent biological experiments were performed for each experiment.

### Structure modelling

The model of an Smc1-Smc3 heterodimer with coiled-coil extensions from the ATPase head domains was created by a combination of the Smc1 homodimer structure (PDB entry 1W1W) and the Smc3:Scc1 complex structure (PDB entry 4UX3) as outlined previously ^30,48^. The model was developed to be consistent with the cross-linking data using the program Chimera (Pettersen et al., 2004, by manual adjustment of the locations of the coiled-coils for each Smc protein, using the Smc3 structure as a template and correcting for the sequence of the Smc1 protein chain. The coiled-coil sections were aligned such that the positions of residues identified by the cross- linking studies were consistent with distances required to make covalent cross-links and such that allowed values for main chain rotation angles were still achieved

## ACKNOWLEDGEMENTS

We are grateful to Hu and Nasmyth lab members for useful discussions, A Donaldson and A Lorenz for critical reading of the manuscript. B Hu was supported by BBSRC (BB/S002537) and the Wellcome Trust (202062/Z/16/Z).

## Author Contributions

A.K. and B.H. designed and conducted experiments and wrote the manuscript. T.T. designed and conducted experiments. N.J.P. performed calibrated ChIP-seq and revised the manuscript. M.V. purified cohesin tetramer/Scc2 and conducted ATPase analysis. C.P. and P.D. conducted experiments. J.B.R conducted protein structure modelling. K.A.N. designed experiments and wrote the manuscript.

