## Supplementary Figures for "Conformational dynamics of cohesin/Scc2 loading complex are regulated by Smc3 acetylation and ATP binding"

Supplementary Figure S1

A

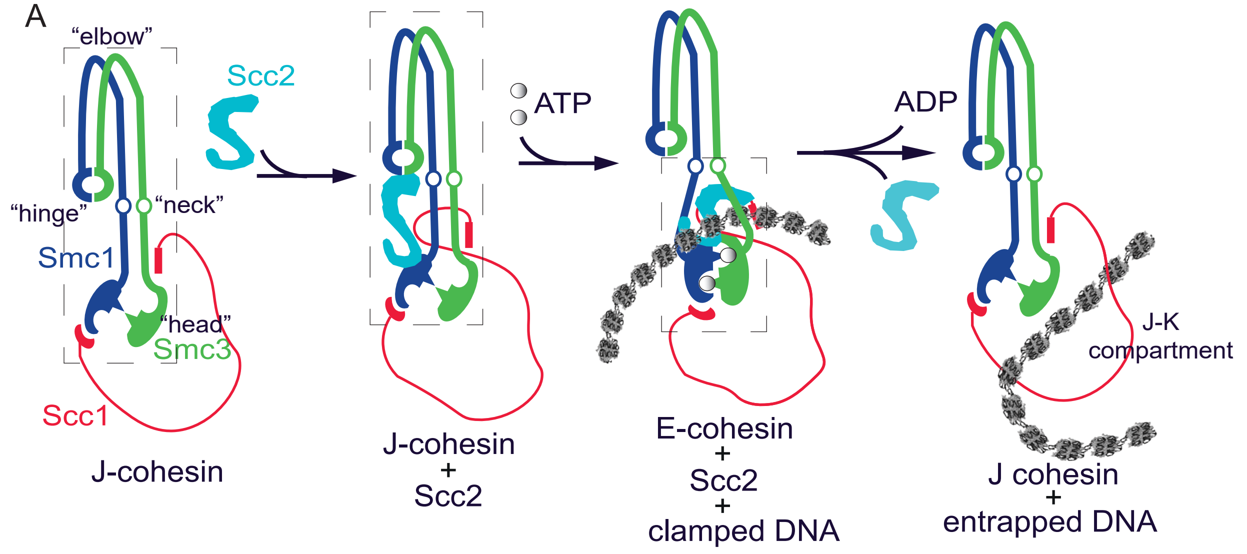


B

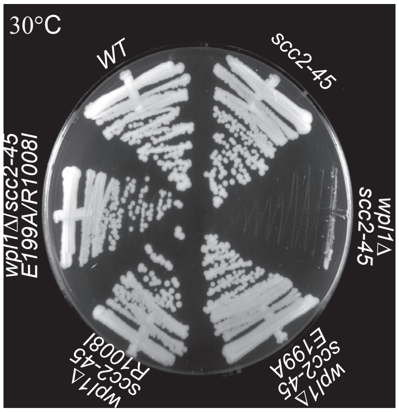


C

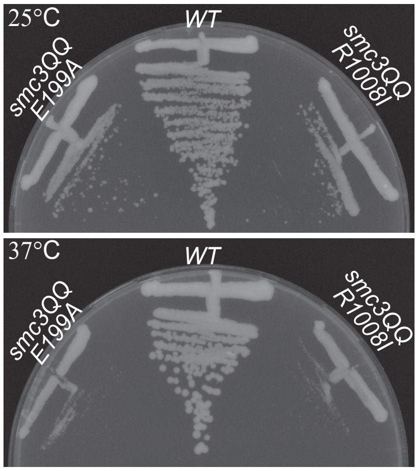

Supplementary Figure S2

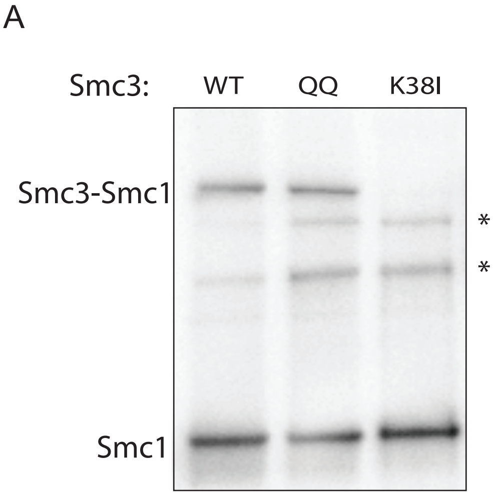


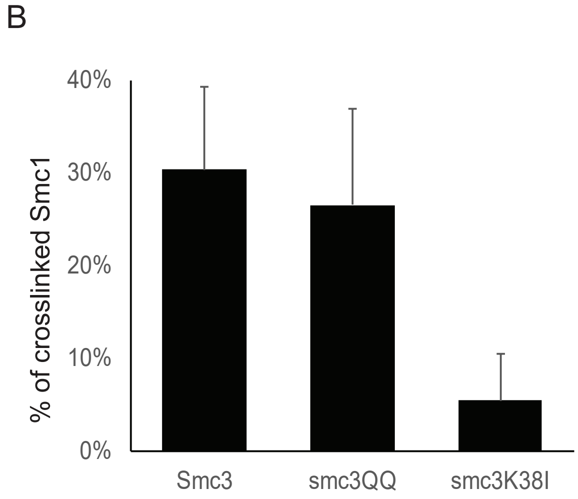

Supplementary Figure S3

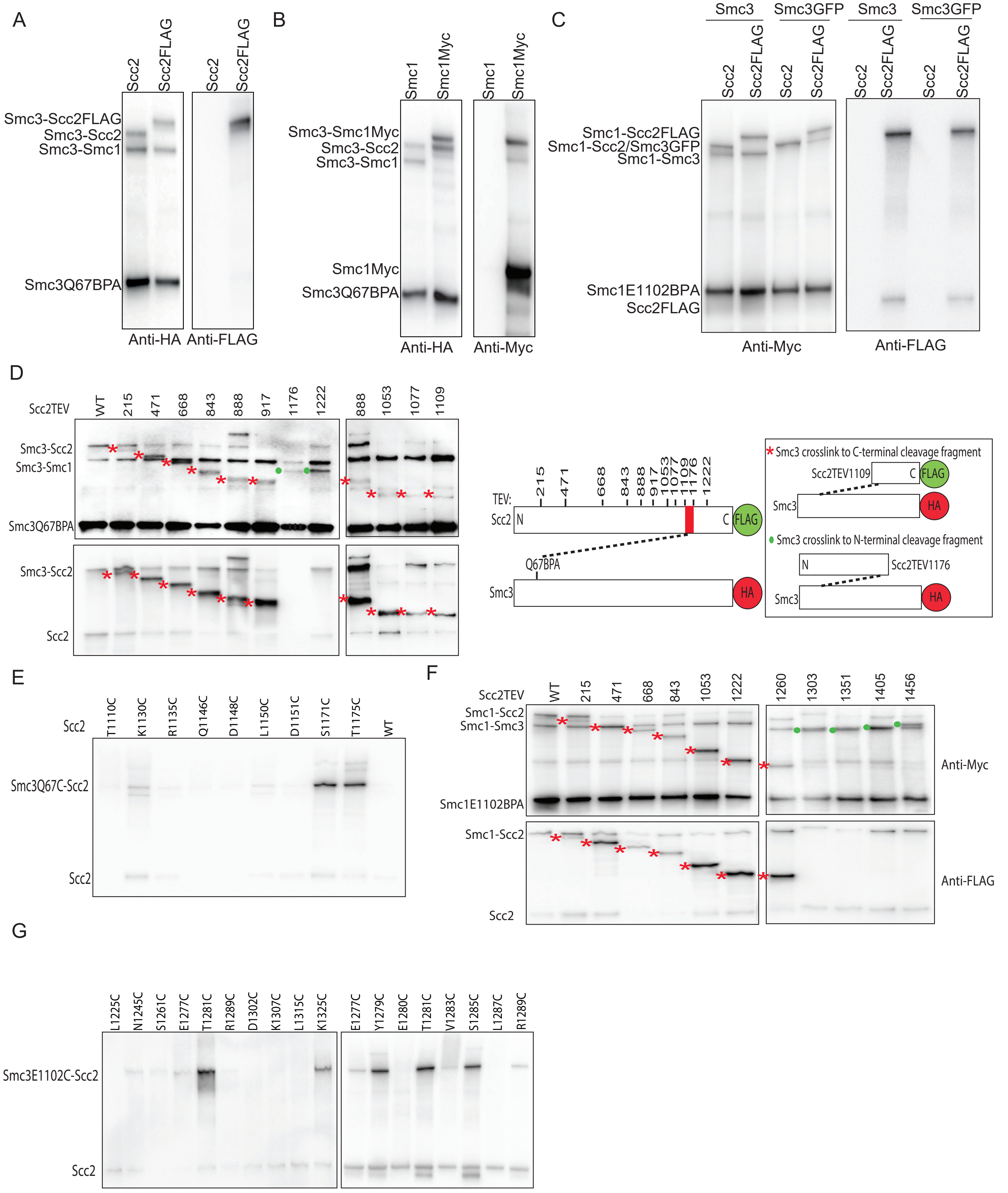

#### Supplementary Figure S4

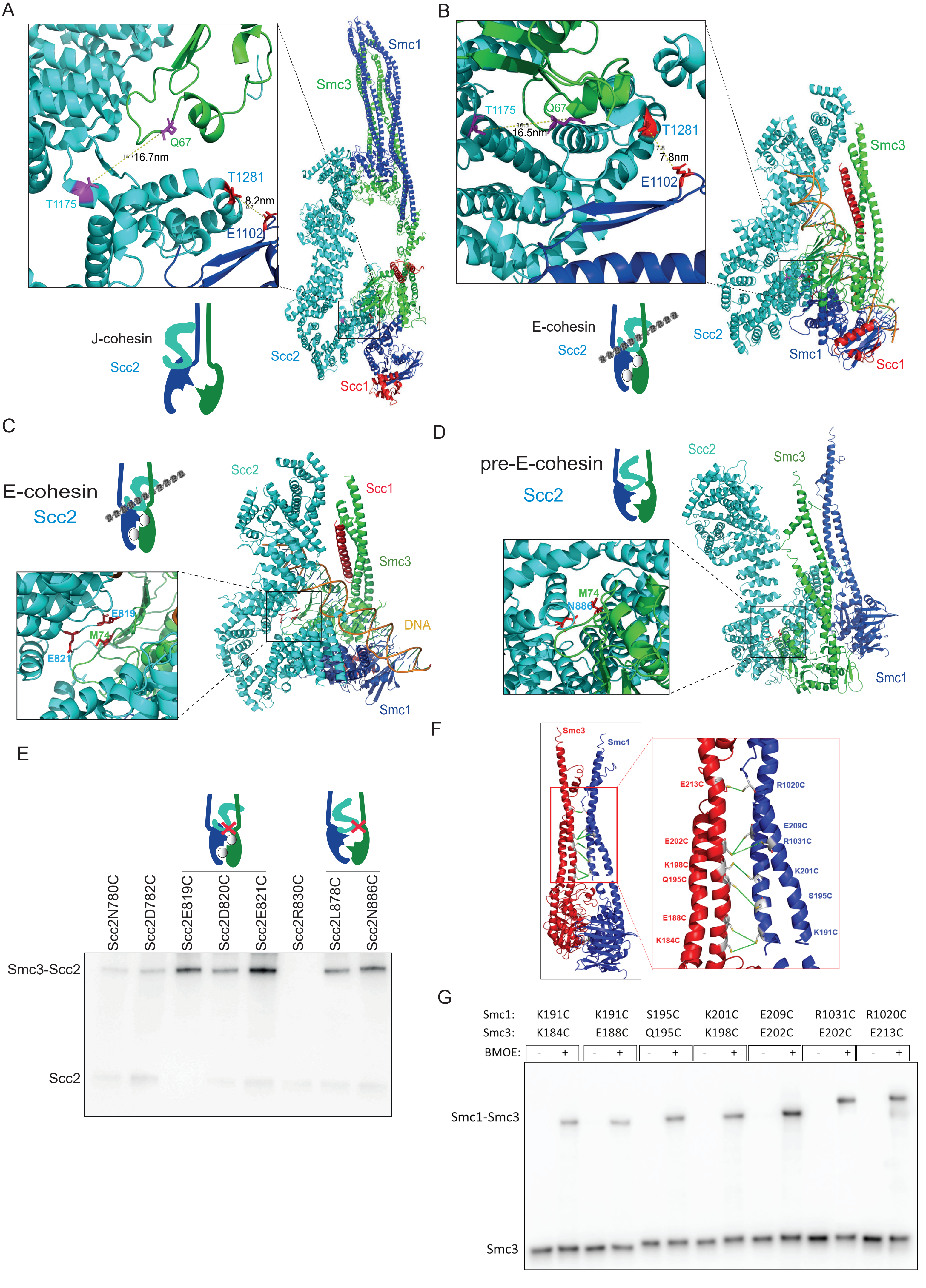

Supplementary Figure S5

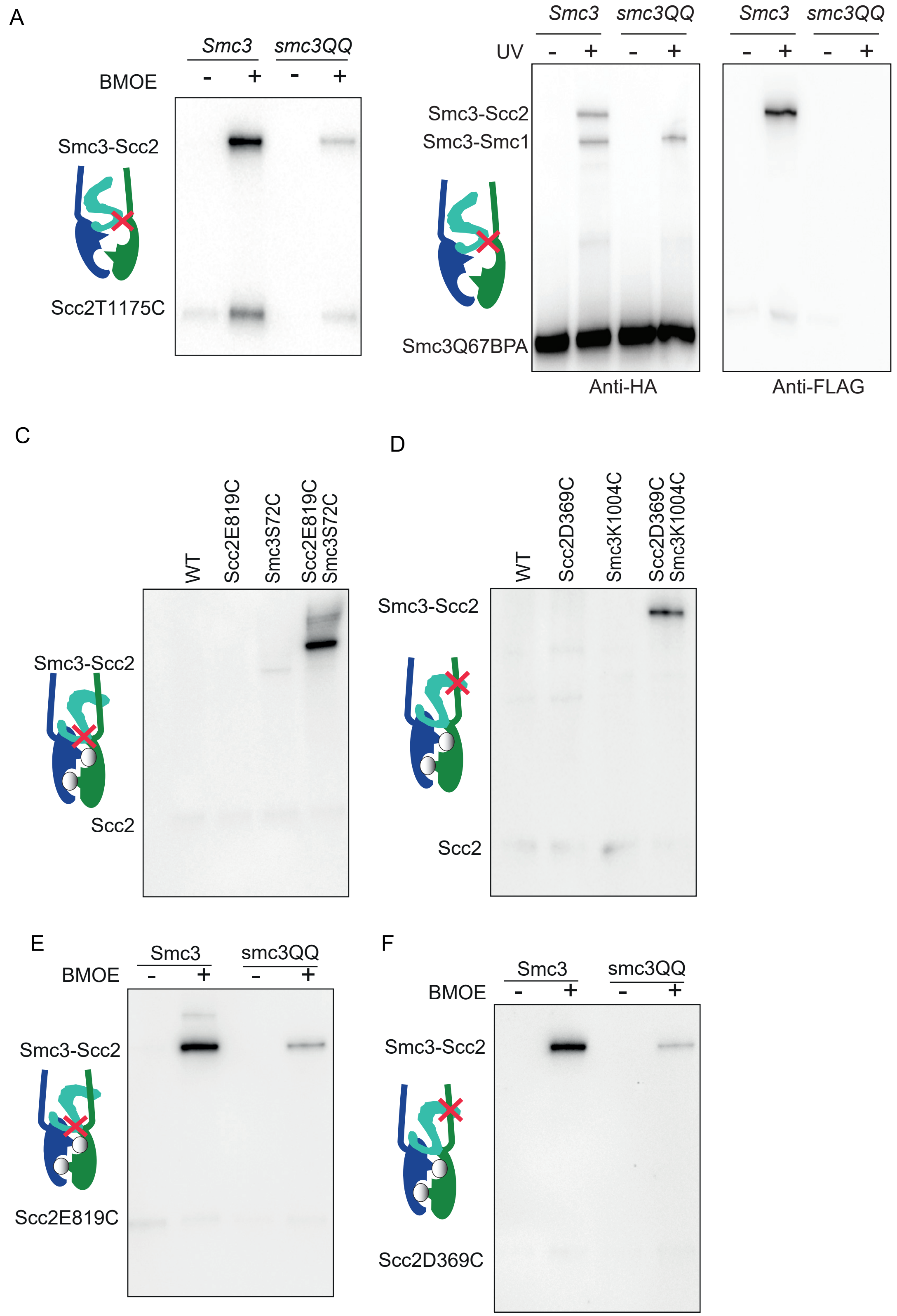

### Supplementary Figure S6

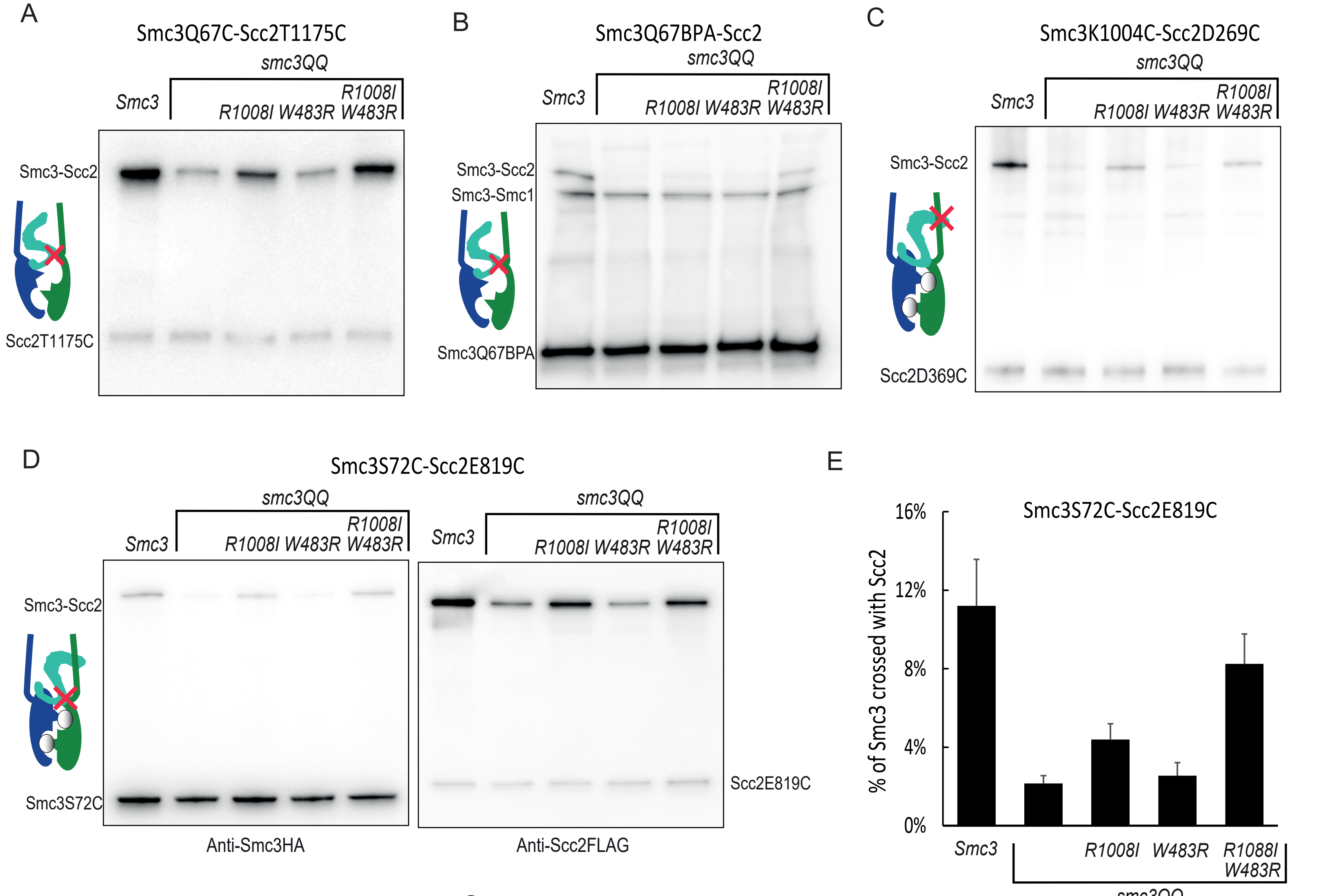


F

| | | |
|---|---|---|
| Q loop | N156D | A157V |
| | K158E | E1124K |
| Signature motif | V1133I | Q1143R |
| | S1176P | |
| ATP binding pocket | F41L | C1183S |
| | K1201E | E1203G |

G

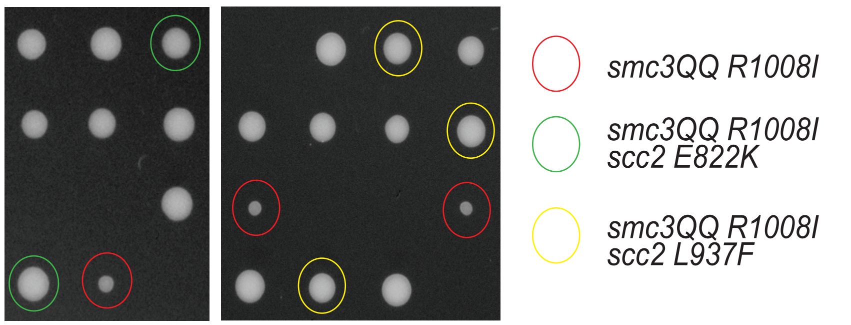


H

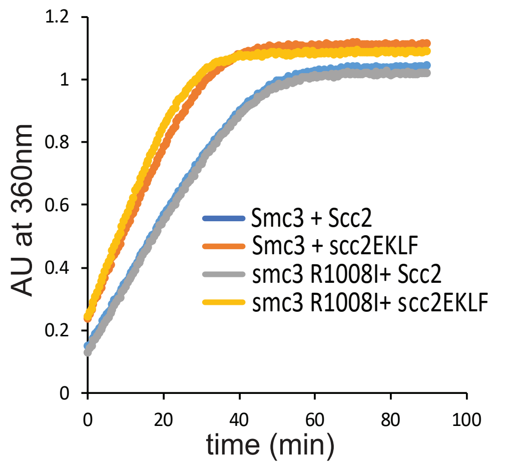

Supplementary Figure S7

A

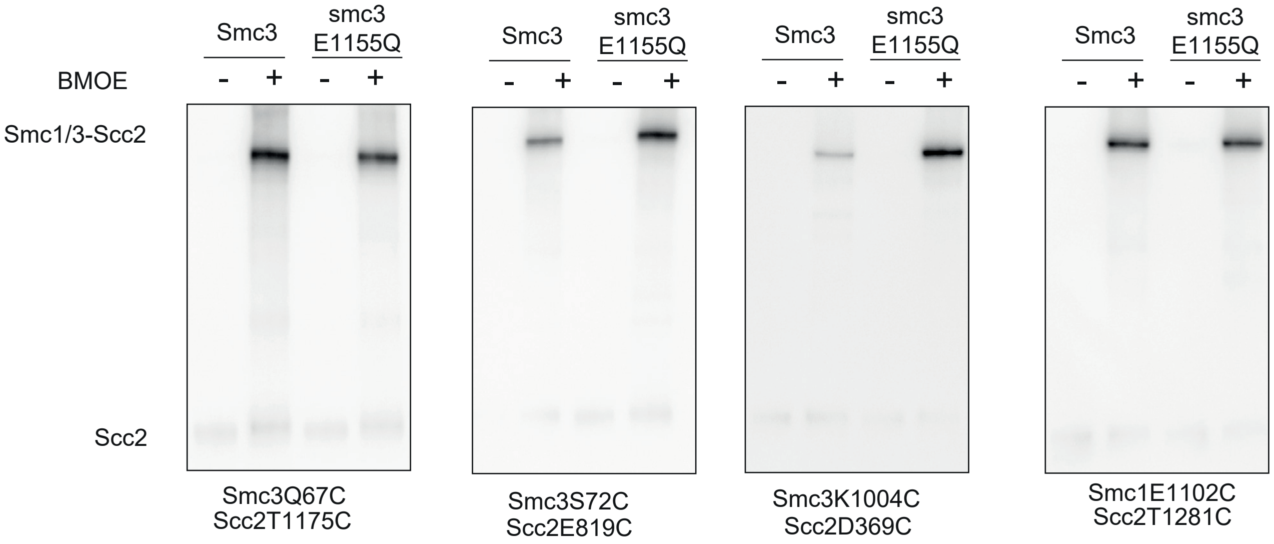
