## Supplemental materials and figure legends for "Conformational dynamics of cohesin/Scc2 loading complex are regulated by Smc3 acetylation and ATP binding"

**Supplementary Fig 1**

(A) Summary of recent discoveries of cohesin loading: cohesin is normally presented with juxtaposed CCs (J-cohesin). The Scc2 forms a complex with J-cohesin through Scc2/Smc1 head interaction. Then, Scc2/cohesin complex is recruited to DNA. In this configuration, cohesin heads are engaged by ATP binding (E-cohesin), Scc2 interacts with both heads, and DNA is clamped in a compartment enclosed by Scc2 and Smc3 CC. After ATP hydrolysis, E-cohesin is transformed into J-cohesin, and DNA is entrapped in the J-K compartment. The squares with dash lines indicate the structures revealed by Cryo-EM analysis.
(B) The growth of indicated strains after incubation at 30°C for two days
(C) The growth of indicated strains after incubation at 25°C or 37°C for two days

**Supplementary Fig 2**

(A) Examine Smc1 by western blot in the crosslink shown in Fig 2C. Asterisks show the positions of non-specific bands.
(B) The percentage of crosslinking efficiency between Smc1 and Smc3 heads was calculated as mean ± SD of 3 independent experiments. The percentage of cross-link efficiency is shown in the graph.

**Supplementary Fig 3**

(A) *In vivo* BPA crosslink of Smc3 Q67BPA with Scc2. Immunoprecipitation of Scc1-PK, Smc3Q67BPA-HA and Scc2-FLAG were detected by Western Blot. Fusing Scc2 with 6xFLAG confirmed the top band as a Smc3-Scc2 crosslink product.
(B) *In vivo* BPA crosslink of Smc3 Q67BPA. Immunoprecipitated on Scc1-PK, separated on gradient gel, and analysed by western blot. Fusing Smc1 with 9xMyc confirmed Smc3 Q67BPA’s crosslink to Smc1.
(C) *In vivo* BPA crosslink of Smc1E1102BPA. Immunoprecipitated on Scc1-PK, separated on gradient gel, and analysed by western blot. Fusing Smc3 with GFP and Scc2 with 6xFLAG revealed the top crosslink band to be Smc1-Scc2 and lower to be Smc1-Smc3.
(D) The Smc3 Q67BPA-Scc2TEV crosslink was digested by TEV protease and cleaved fragments were examined by Western Blot. The BOME crosslink of Smc3 Q67C with indicated cysteine substituted Scc2 as shown in Fig 2B. Scc2 was detected using anti-Flag antibody.
(E) The BOME crosslink of Smc3 Q67C with indicated cysteine substituted Scc2. Scc2 was detected using anti-Flag antibody.
(F) The Smc1 E1102BPA-Scc2TEV crosslink was digested by TEV protease and cleaved fragments were examined by Western Blot.
(G) The BOME crosslink of Smc1 E1102C with indicated cysteine substituted Scc2. Scc2 was detected using anti-Flag antibody.

**Supplementary Fig 4**

(A) The indicated distance between Smc3 Q67 and Scc2 or Smc1 E1102 and Scc2T1281 in the Scc2/J-cohesin complex
(B) The indicated distance between Smc3 Q67 and Scc2 or Smc1 E1102 and Scc2T1281 in the Scc2/E-cohesin complex.
(C) The close proximity of Smc3 M74 with Scc2 E819 and Scc2 E821 in the Scc2/E-cohesin complex.
(D) The close proximity of Smc3 M74 with Scc2 N888 in Scc2/pre-E-cohesin complex.
(E) Scc2 was detected by Western Blot from the crosslink shown Fig 4B.
(F) Structural model of juxtaposed cohesin CCs.
(G) *In vivo* BOME crosslink of indicated cysteine pairs on cohesin CCs.

**Supplementary Fig 5**

(A) The BOME crosslink of indicated cysteine substituted Scc2 as shown in Fig 5A. Scc2 was detected by Western Blot.
(B) *In vivo* BPA crosslink of Smc3 or smc3QQ Q67BPA with Scc2. After the immunoprecipitation of Scc1-PK, Smc3 Q67BPA-HA and Scc2-FLAG were detected by Western Blot.
(C) Scc2 was detected by Western Blot from the crosslink shown in Fig 5D.
(D) Scc2 was detected by Western Blot from the crosslink shown in Fig 5E.
(E) Scc2 was detected by Western Blot from the crosslink shown in Fig 5F.
(F) Scc2 was detected by Western Blot from the crosslink shown in Fig 5H.

**Supplementary Fig 6**

(A) Scc2 was detected by Western Blot from the crosslinks shown in Fig 7B
(B) *In vivo* BPA crosslink revealed the effects of Smc3 R1008 and W483R on the interface between *smc3QQ Q67* and Scc2.
(C) Scc2 was detected by Western Blot from the crosslinks shown in Fig 7D
(D) Scc2 S72C and *smc3QQ* E819C were crosslinked *in vivo* in the presence of R1008I and W483R or both using BMOE. Immunoprecipitated on Scc1-PK, separated on gradient gel, and analysed by Western Blot.
(E) The percentage of crosslinking products in panel D was calculated as mean+SD of 3 independent experiments. A combination of Smc3 R1008I and W384R increased the crosslink of smc3QQ with Scc2 to about 70% of wild type.
(F) The distributions of second suppressor mutations on Smc3 head
(G) Either Scc2 single mutation, E822K (green circles) or L937F (yellow circles) promotes cell proliferation of smc3QQ R1008I mutant (red circles).
(H) Recombinant tetramer of cohesin, Smc3 or Smc3 R1008I, Smc1, Scc1 and Scc3 was incubated with Scc2 or Scc2 E822K L937F. ATP was added to initiate the reaction and reaction rate was measured as the change in absorption at 360nm over time.

**Supplementary Fig 7**

(A) Scc2 was detected by Western Blot from the crosslinks shown in Fig 7A.
