## Supplementary material for "Conformational dynamics of cohesin/Scc2 loading complex are regulated by Smc3 acetylation and ATP binding": Table S1

Table S1 Cysteine pairs used in this study

| **Cysteine pair** | **Interface** | **Configuration** |
| --- | --- | --- |
| Smc3 S1043C  Scc1 C56 | Smc3 CC/N-Scc1 | Interacted Smc3 CC/N-Scc1 |
| Smc1 N1192C  Smc3 R1222C | Cohesin heads | Engaged heads |
| Smc1 R1031C  Smc3 E202C | Cohesin CCs | Juxtaposed CCs |
| Smc1 E1102C  Scc2 T1281C | Scc2/Smc1 head | Scc2/J-cohesin  Scc2/pre-E-cohesin  Scc2/E-cohesin |
| Smc3 Q67C  Scc2 T1175C | Scc2/Smc3 head | Scc2/pre-E-cohesin |
| Smc3 M74C  Scc2 N886C |  |  |
| Smc3 S72C  Scc2 E819C | Scc2/Smc3 head | Scc2/E-cohesin |
| Smc3 M74C  Scc2 E821C |  |  |
| Smc3 K1004C  Scc2 D369C | Scc2/Smc3 CC |  |
