## Supplementary material for "Conformational dynamics of cohesin/Scc2 loading complex are regulated by Smc3 acetylation and ATP binding": Table S2

Table S1 yeast strains used in this study

| **Figures** | **Genotype** | **Origin** | **Strains** |
| --- | --- | --- | --- |
|  | *All the S. cerevisiae strains derive from W303:*  *ade2-1, trp1-1, can1-100, leu2-3,112,his3-11,15, ura3, GAL, psi+* | Nasmyth’ lab | K699 |
| **Figure 1 and S1** | | | |
| 1A | *Mat a/alpha, Δsmc3::HIS3/Smc3, ura3::smc3(K112Q_K113Q)::URA3/ura3* | This study | B1356 |
|  | *Mat a/alpha, Δsmc3::HIS3/Smc3, ura3::smc3(K112Q_K113Q_E199A)::URA3/ura3* | This study | B1931 |
|  | *Mat a/alpha, Δsmc3::HIS3/Smc3, ura3::smc3(K112Q_K113Q_R1008I)::URA3/ura3* | This study | B1335 |
| 1C | *Mat a, trp1::SMC3-PK6::TRP1* | This study | K17407 |
|  | *Mat a, trp1::smc3(K112Q_K113Q)-PK6::TRP1* | This study | K22703 |
|  | *Mat a, trp1::smc3(K112Q_K113Q_R1008I)-PK6::TRP1* | This study | K22705 |
| 1E | *Mat a/alpha, Δsmc3::HIS3/SMC3,*  *ura3::smc3(K112Q_K113Q_R1008I_W483R)::URA3/ura3,*  *SCC1-PK9::KANMX/scc1* | This study | B1471 |
| 1F | *MATa, SCC1-PK9::KANMX* | This study | B910 |
|  | *Mat a, SCC1-PK9::KANMX, Δsmc3::HIS3, ura3::smc3(K112Q_K113Q_R1008I)::URA3* | This study | B2269 |
|  | *Mat a, SCC1-PK9::KANMX, Δsmc3::HIS3, ura3::smc3(K112Q_K113Q_R1008I_W483R)::URA3* | This study | B2851 |
| 1G | *MAT a/alpha, Δeco1::KANMX6/Eco1, smc3(W483R_R1008I)::HIS3/SMC3* | This study | B2902 |
|  | *MAT a/alpha, Δeco1::KANMX6/Eco1, smc3(K112Q_K113Q_W483R_R1008I)::His3/SMC3* | This study | B2903 |
| 1H | *ura3::smc3(S1043C)-PK6::URA3* | This study | B484 |
|  | *ura3:smc3(K112Q, K113Q, S1043C)-PK6::URA3* | This study | B482 |
|  | *ura3::smc3(K112Q, K113Q, W483R, R1008I, S1043C)-PK6::URA3* | This study | B4020 |
| S1B | *MAT a* | Nasmyth | K699 |
|  | *MAT a, scc2-45::NATMX (L545P D575G)* | This study | B443 |
|  | *MAT a, scc2-45::NATMX (L545P D575G), Δwpl1::HPHMX4* | This study | B1286 |
|  | *MAT a, scc2-45::NATMX (L545P D575G), Δwpl1::HPHMX4, smc3(R1008I)::HIS3* | This study | B1284 |
|  | *MAT a, scc2-45::NATMX (L545P D575G), Δwpl1::HPHMX4, smc3(E199A)::HIS3* | This study | K22222 |
|  | *MAT a, scc2-45::NATMX (L545P D575G), Δwpl1::HPHMX4, smc3(E199A_R1008I)::HIS3* | This study | K22224 |
| S1C | *MAT a* | Nasmyth | K699 |
|  | *Mat a/alpha, Δsmc3::HIS3/SMC3, ura3::smc3(K112Q_K113Q_E199A)::URA3/ura3* | This study | B1931 |
|  | *Mat a/alpha, Δsmc3::HIS3/SMC3, ura3::smc3(K112Q_K113Q_R1008I)::URA3/ura3* | This study | B1931 |
|  | *Mat a, Δsmc3::HIS3, ura3::smc3(K112Q_K113Q_E199A)::URA3* | This study | Dissected from B1931 |
|  | *Mat a, Δsmc3::HIS3, ura3::smc3(K112Q_K113Q_R1008I)::URA3* | This study | Dissected from B1931 |
| **Figure 2 and S2** | | | |
| 2C and S2A | *Mat a, smc1::NATMX4, SCC1-PK9::KANMX, Δmet15::smc1N1192C-myc9, ura3::smc3R1222C-HA3::URA3* | This study | B2936 |
|  | *Mat a, smc1::NATMX4, SCC1-PK9::KANMX, Δmet15::smc1N1192C-myc9, ura3::smc3K112Q_K113Q_R1222C-HA3::URA3* | This study | B2937 |
|  | *Mat a, Δsmc1::NATMX4, SCC1-PK9::KanMX, Δmet15::smc1N1192C-myc9, ura3::smc3K38I_R1222C-HA3::URA3* | This study | B3046 |
| 2E | *Mat a, SCC2_6xHis-FLAG6::KANMX* | This study | B1449 |
|  | *Mat a, SCC2_6xHis-FLAG6::KANMX, ura3::SMC3-PK6::URA3* | This study | B3026 |
|  | *Mat a, SCC2_6xHis-FLAG6::KANMX, ura3::smc3(K112Q_K113Q)-PK6::URA3* | This study | B3027 |
| **Figure 3 and S3** | | | |
| 3B | *MATa, SCC1-PK9::KANMX, SCC2_6xHis-FLAG6::KANMX, pBH760 (smc3-L111TAG-HA3 ), pBH61 (tRNATyr/Bpa synthetase)* | This study | B1852 |
|  | *MATa, SCC1-PK9::KANMX, SCC2_6xHis-FLAG6::KANMX, pBH761 (smc3-Q117TAG-HA3), pBH61 (tRNATyr/Bpa synthetase)* | This study | B1855 |
|  | *MATa, SCC1-PK9::KANMX, SCC2_6xHis-FLAG6::KANMX, pBH758 (smc3-K57TAG-HA3 ), pBH61 (tRNATyr/Bpa synthetase)* | This study | B1850 |
|  | *MATa, SCC1-PK9::KANMX, SCC2_6xHis_FLAG6::KANMX, pBH796 (smc3-R58TAG-HA3), pBH61 (tRNATyr/Bpa synthetase)* | This study | B1900 |
|  | *MATa, SCC1-PK9::KANMX, SCC2_6xHis_FLAG6::KANMX, pBH773 (smc3-R61TAG-HA3 ), pBH61 (tRNATyr/Bpa synthetase)* | This study | B1848 |
|  | *MATa, SCC1-PK9::KANMX, SCC2_6xHis_FLAG6::KANMX, pBH759 (smc3-M74TAG-HA3 ), pBH61 (tRNATyr/Bpa synthetase)* | This study | B1854 |
|  | *MATa, SCC1-PK9::KANMX, SCC2_6xHis_FLAG6::KANMX*  *pBH787 (Smc3-H66TAG-HA3), pBH61 (tRNATyr/Bpa synthetase)* | This study | B1878 |
|  | *MATa, SCC1-PK9::KANMX, SCC2_6xHis_FLAG6::KANMX, pBH585 (smc3-Q67TAG-HA3 ), pBH61 (tRNATyr/Bpa synthetase)* | This study | B1505 |
| 3D | *MATa, SCC1-PK9::KANMX, SCC2_6xHis_FLAG6::KANMX*  *pBH1059 (smc1D1069TAG-myc9), pBH61 (tRNATyr/Bpa synthetase)* | This study | B3201 |
|  | *MATa, SCC1-PK9::KANMX, SCC2_6xHis_FLAG6::KANMX*  *pBH1060 (smc1D1073TAG-myc9), pBH61 (tRNATyr/Bpa synthetase)* | This study | B3202 |
|  | *MATa, SCC1-PK9::KANMX, SCC2_6xHis_FLAG6::KANMX*  *pBH1061 (smc1D1076TAG-myc9), pBH61 (tRNATyr/Bpa synthetase)* | This study | B3203 |
|  | *MATa, SCC1-PK9::KANMX, SCC2_6xHis_FLAG6::KANMX*  *pBH1062 (smc1R1080TAG-myc9), pBH61 (tRNATyr/Bpa synthetase)* | This study | B3204 |
|  | *MATa, SCC1-PK9::KANMX, SCC2_6xHis_FLAG6::KANMX*  *pBH1055 (smc1T1100TAG-myc9), pBH61 (tRNATyr/Bpa synthetase)* | This study | B3199 |
|  | *MATa, SCC1-PK9::KANMX, SCC2_6xHis_FLAG6::KANMX, pBH1056 (smc1E1102TAG-Myc9), pBH61 (tRNATyr/Bpa synthetase)* | This study | B2814 |
|  | *MATa, Scc1-PK9::KANMX, SCC2_6xHis_FLAG6::KANMX*  *pBH1057 (smc1K1113TAG-Myc9), pBH61 (tRNATyr/Bpa synthetase)* | This study | B2815 |
|  | *MATa, SCC1-PK9::KANMX, SCC2_6xHis_FLAG6::KANMX*  *pBH1058 (smc1T1117TAG-myc9), pBH61 (tRNATyr/Bpa synthetase)* | This study | B3200 |
|  | *MATa, SCC1-PK9::KANMX, SCC2_6xHis_FLAG6::KANMX, pBH1047 (smc1L1120TAG-Myc9 ), pBH61 (tRNATyr/Bpa synthetase)* | This study | B3193 |
|  | *MATa, SCC1-PK9::KANMX, SCC2_6xHis_FLAG6::KANMX, pBH1048 (smc1R1122TAG-Myc9 ), pBH61 (tRNATyr/Bpa synthetase)* | This study | B2811 |
|  | *MATa, SCC1-PK9::KANMX, SCC2_6xHis_FLAG6::KANMX*  *pBH1049 (smc1F1123TAG-myc9), pBH61 (tRNATyr/Bpa synthetase)* | This study | B3196 |
|  | *MATa, SCC1-PK9::KANMX, SCC2_6xHis_FLAG6::KANMX, pBH1050 (smc1K1124TAG-Myc9 ), pBH61 (tRNATyr/Bpa synthetase)* | This study | B2812 |
|  | *MATa, SCC1-PK9::KANMX, SCC2_6xHis_FLAG6::KANMX, pBH1051 (smc1E1127TAG-Myc9 ), pBH61 (tRNATyr/Bpa synthetase)* | This study | B2813 |
|  | *MATa, SCC1-PK9::KANMX, SCC2_6xHis_FLAG6::KANMX*  *pBH1052 (smc1Y1128TAG-myc9), pBH61 (tRNATyr/Bpa synthetase)* | This study | B3197 |
| 3E | *MATa, SCC1-PK9::KANMX, Δscc2::NATMX4, smc3HA6::HIS3, Δlys2:: SCC2_6xHis_FLAG6* | This study | B3161 |
|  | *MATa, SCC1-PK9::KANMX, Δscc2::NATMX4, smc3HA6::HIS3, Δlys2::scc2T1175C_HIS6_FLAG6* | This study | B3176 |
|  | *MATa, SCC1-PK9::KANMX, Δscc2::NATMX4, smc3_Q67C_HA::His3, Δlys2:: SCC2_6xHis_FLAG6* | This study | B3054 |
|  | *MATa, , SCC1-PK9::KANMX, Δscc22::NATMX4, smc3_Q67C_HA::His3, Δlys2::scc2T1175C_HIS6_FLAG6* | This study | B3064 |
| 3F | *MATa,smc1-myc9:: kiTRP1, SCC1-PK9::KANMX, Δscc2::NATMX4, Δlys2:: SCC2_6xHis_FLAG6* | This study | B3115 |
|  | *MATa, smc1-myc9::kiTRP1, SCC1-PK9::KANMX, Δscc2::NATMX4, Δlys2:: scc2T1281C_HIS6_FLAG6,* | This study | B3110 |
|  | *MATa, SCC1-PK9::KANMX, Δscc2::NATMX4, smc1E1102Cmyc9:: kiTRP1, Δlys2:: SCC2_6xHis_FLAG6* | This study | B3097 |
|  | *MATa, SCC1-PK9::KANMX, Δscc2::NATMX4, smc1E1102Cmyc9:: kiTRP1, Δlys2::scc2T1281C_HIS6_FLAG6* | This study | B3043 |
| 3G | *MATa, , SCC1-PK9::KANMX, Δscc22::NATMX4, smc3_Q67C_HA::His3, Δlys2::scc2T1175C_HIS6_FLAG6* | This study | B3064 |
|  | *MATa, SCC1-PK9::KANMX, Δscc2::NATMX4, smc1E1102Cmyc9:: kiTRP1, Δlys2::scc2T1281C_HIS6_FLAG6* | This study | B3043 |
|  | *MATa, SCC1-PK9::KANMX, Δscc2::NATMX4, smc3Q67C_HA6::spHis5, Δlys2::Scc2T1281C_HIS6_FLAG6* | This study | B3174 |
|  | *MATa, SCC1-PK9::KANMX, ΔSCC2::NATMX4, smc1E1102Cmyc9:: kiTRP1, Δlys2:: scc2T1175C_HIS6_FLAG6* | This study | B3172 |
|  | *MATa, SCC1-PK9::KANMX, Δscc2::NATMX4, smc3Q67C_HA6::spHis5, smc1E1102Cmyc9:: kiTRP1, Δlys2:: scc2_T1175C_T1281C_HIS6_FLAG6* | This study | B3194 |
| S3A and S3B | *Mat a, SCC1-PK9::KanMX, pBH585 (smc3-Q67TAG-HA3), pBH61 (tRNATyr/Bpa synthetase)* | This study | B1613 |
|  | *Mat a, SCC1-PK9::KanMX, SCC2_6xHis_FLAG6::KANMX, pBH585 (smc3-Q67TAG-HA3), pBH61 (tRNATyr/Bpa synthetase)* | This study | B1505 |
|  | *Mat a, SCC1-PK9::KanMX, Δsmc1::NatMX4, URA3::Smc1-Myc9::URA3, pBH585 (Smc3-Q67TAG-HA3), pBH61 (tRNATyr/Bpa synthetase)* | This study | B1614 |
| S3C | *MATa, SCC1-PK9::KANMX, pBH1056 (smc1E1102TAG-Myc9), pBH61 (tRNATyr/Bpa synthetase)* | This study | B2797 |
|  | *MATa, SCC1-PK9::KANMX, SCC2_6xHis_FLAG6::KANMX, pBH1056 (smc1E1102TAG-Myc9), pBH61 (*tRNA^Tyr^/Bpa synthetase*)* | This study | B2814 |
|  | *Mat a, SMC3-GFP::URA3, SCC1-PK9::KANMX, pBH1056(smc1E1102TAG-Myc9 in YEplac181), pBH61 (tRNATyr/Bpa synthetase)* | This study | B2980 |
|  | *Mat a, SMC3-GFP::URA3, SCC1-PK9::KANMX, SCC2_6xHis_FLAG6::KANMX, pBH1056(smc1E1102TAG-Myc9 in YEplac181), pBH61 (tRNATyr/Bpa synthetase)* | This study | B3028 |
| S3D | *MATa, SCC1-PK9::KANMX, SCC2_6xHis_FLAG6::KANMX, pBH1056 (smc1E1102TAG-Myc9), pBH61 (tRNATyr/Bpa synthetase)* | This study | B2814 |
|  | *MAT a, SCC1-PK9::KANMX, scc2TEV215_6xHis_FLAG6:: KANMX, pBH61 (tRNATyr/Bpa synthetase), pBH585 (smc3-Q67TAG-HA3)* | This study | B2171 |
|  | *MAT a, SCC1-PK9::KANMX, scc2TEV471_6xHis_FLAG6:: KANMX, pBH61 (tRNATyr/Bpa synthetase), pBH585 (smc3-Q67TAG-HA3)* | This study | B2172 |
|  | *MAT a, SCC1-PK9::KANMX, scc2TEV668_6xHis_FLAG6:: KANMX, pBH61 (tRNATyr/Bpa synthetase), pBH585 (smc3-Q67TAG-HA3)* | This study | B2145 |
|  | *MAT a, SCC1-PK9::KANMX, scc2TEV843_6xHis_FLAG6:: KANMX, pBH61 (tRNATyr/Bpa synthetase), pBH585 (smc3-Q67TAG-HA3)* | This study | B2149 |
|  | *MAT a, SCC1-PK9::KANMX, scc2TEV888_6xHis_FLAG6:: KANMX, pBH61 (tRNATyr/Bpa synthetase), pBH585 (smc3-Q67TAG-HA3)* | This study | B2152 |
|  | *MAT a, SCC1-PK9::KANMX, scc2TEV917_6xHis_FLAG6:: KANMX, pBH61 (tRNATyr/Bpa synthetase), pBH585 (smc3-Q67TAG-HA3)* | This study | B2147 |
|  | *MAT a, SCC1-PK9::KANMX, scc2TEV1176_6xHis_FLAG6:: KANMX, pBH61 (tRNATyr/Bpa synthetase), pBH585 (smc3-Q67TAG-HA3)* | This study | B2170 |
|  | *MAT a, SCC1-PK9::KANMX, scc2TEV1222_6xHis_FLAG6:: KANMX, pBH61 (tRNATyr/Bpa synthetase), pBH585 (smc3-Q67TAG-HA3)* | This study | B2151 |
|  | *MAT alpha, SCC1-PK9::KANMX, scc2TEV1053_6xHis_FLAG6:: KANMX, pBH61 (tRNATyr/Bpa synthetase), pBH585 (smc3-Q67TAG-HA3)* | This study | B2767 |
|  | *MAT a, SCC1-PK9::KANMX, scc2TEV1077_6xHis_FLAG6:: KANMX, pBH61 (tRNATyr/Bpa synthetase), pBH585 (smc3-Q67TAG-HA3)* | This study | B2768 |
|  | *MAT a SCC1-PK9::KANMX, scc2TEV1109_6xHis_FLAG6:: KANMX, pBH61 (tRNATyr/Bpa synthetase), pBH585 (smc3-Q67TAG-HA3)* | This study | B2769 |
| S3E | *MATa, SCC1-PK9::KANMX, Δscc2::NATMX4*  *smc3_Q67C_HA6::spHis5, Δlys2:: scc2T1110C_HIS6_FLAG6* | This study | B3050 |
|  | *MATa, SCC1-PK9::KANMX, Δscc2::NATMX4*  *smc3_Q67C_HA6::spHis5, Δlys2:: scc2K11300C_HIS6_FLAG6* | This study | B3080 |
|  | *MATa, SCC1-PK9::KANMX, Δscc2::NATMX4*  *smc3_Q67C_HA6::spHis5, Δlys2:: scc2R1135C_HIS6_FLAG6* | This study | B3062 |
|  | *MATa, SCC1-PK9::KANMX, Δscc2::NATMX4*  *smc3_Q67C_HA6::spHis5, Δlys2:: scc2Q1146C_HIS6_FLAG6* | This study | B3051 |
|  | *MATa, SCC1-PK9::KANMX, Δscc2::NATMX4*  *smc3_Q67C_HA6::spHis5, Δlys2:: scc2D1148C_HIS6_FLAG6* | This study | B3061 |
|  | *MATa, SCC1-PK9::KANMX, Δscc2::NATMX4*  *smc3_Q67C_HA6::spHis5, Δlys2:: scc2L1150C_HIS6_FLAG6* | This study | B3052 |
|  | *MATa, SCC1-PK9::KANMX, Δscc2::NATMX4*  *smc3_Q67C_HA6::spHis5, Δlys2:: scc2D1151C_HIS6_FLAG6* | This study | B3063 |
|  | *MATa, SCC1-PK9::KANMX, Δscc2::NATMX4*  *smc3_Q67C_HA6::spHis5, Δlys2:: scc2S1171C_HIS6_FLAG6* | This study | B3053 |
|  | *MATa, SCC1-PK9::KANMX, Δscc2::NATMX4*  *smc3_Q67C_HA6::spHis5, Δlys2:: scc2T1175C_HIS6_FLAG6* | This study | B3064 |
|  | *MATa, SCC1-PK9::KANMX, Δscc2::NATMX4*  *smc3_Q67C_HA6::spHis5, Δlys2::SCC2_HIS6_FLAG6* | This study | B3054 |
| S3F | *MATa, SCC1-PK9::KANMX, SCC2_6xHis_FLAG6::KANMX, pBH1056 (smc1E1102TAG-Myc9), pBH61 (tRNATyr/Bpa synthetase)* | This study | B2814 |
|  | *Mat a, SCC1-PK9::KANMX, scc2TEV215_6xHis_FLAG6:: KANMX, pBH1056(smc1E1102TAG-Myc9 in YEplac181), pBH61 (tRNATyr/Bpa synthetase)* | This study | B2906 |
|  | *Mat a, SCC1-PK9::KANMX, scc2TEV471_6xHis_FLAG6:: KANMX, pBH1056(smc1E1102TAG-Myc9 in YEplac181), pBH61 (tRNATyr/Bpa synthetase)* | This study | B2907 |
|  | *Mat a, SCC1-PK9::KANMX, scc2TEV668_6xHis_FLAG6:: KANMX, pBH1056(smc1E1102TAG-Myc9 in YEplac181), pBH61 (tRNATyr/Bpa synthetase)* | This study | B2908 |
|  | *Mat a, SCC1-PK9::KANMX, scc2TEV843_6xHis_FLAG6:: KANMX, pBH1056(smc1E1102TAG-Myc9 in YEplac181), pBH61 (tRNATyr/Bpa synthetase)* | This study | B2909 |
|  | *Mat a, SCC1-PK9::KANMX, scc2TEV1053_6xHis_FLAG6:: KANMX, pBH1056(smc1E1102TAG-Myc9 in YEplac181), pBH61 (tRNATyr/Bpa synthetase)* | This study | B2911 |
|  | *Mat a, SCC1-PK9::KANMX, scc2TEV1222_6xHis_FLAG6:: KANMX, pBH1056(smc1E1102TAG-Myc9 in YEplac181), pBH61 (tRNATyr/Bpa synthetase)* | This study | B2910 |
|  | *Mat a, SCC1-PK9::KANMX, scc2TEV1260_6xHis_FLAG6:: KANMX, pBH1056(smc1E1102TAG-Myc9 in YEplac181), pBH61 (tRNATyr/Bpa synthetase)* | This study | B3017 |
|  | *Mat a, SCC1-PK9::KANMX, scc2TEV1303_6xHis_FLAG6:: KANMX, pBH1056(smc1E1102TAG-Myc9 in YEplac181), pBH61 (tRNATyr/Bpa synthetase)* | This study | B3019 |
|  | *Mat a, SCC1-PK9::KANMX, scc2TEV1351_6xHis_FLAG6:: KANMX, pBH1056(smc1E1102TAG-Myc9 in YEplac181), pBH61 (tRNATyr/Bpa synthetase)* | This study | B3015 |
|  | *Mat a, SCC1-PK9::KANMX, scc2TEV1405_6xHis_FLAG6:: KANMX, pBH1056(smc1E1102TAG-Myc9 in YEplac181), pBH61 (tRNATyr/Bpa synthetase)* | This study | B3016 |
|  | *Mat a, SCC1-PK9::KANMX, scc2TEV1456_6xHis_FLAG6:: KANMX, pBH1056(smc1E1102TAG-Myc9 in YEplac181), pBH61 (tRNATyr/Bpa synthetase)* | This study | B3018 |
| S3G | *MATa, SCC1-PK9::KANMX, scc2::NATMX4*  *smc1E1102Cmyc9:: kiTRP1, Δlys2:: scc2L1225C_HIS6_FLAG6* | This study | B3034 |
|  | *MATa, SCC1-PK9::KANMX, scc2::NATMX4*  *smc1E1102Cmyc9:: kiTRP1, Δlys2:: scc2N1245C_HIS6_FLAG6* | This study | B3035 |
|  | *MATa, SCC1-PK9::KANMX, scc2::NATMX4, smc1E1102Cmyc9:: kiTRP1, Δlys2:: scc2L1261C_HIS6_FLAG6* | This study | B3036 |
|  | *MATa, SCC1-PK9::KANMX, scc2::NATMX4*  *smc1E1102Cmyc9:: kiTRP1, Δlys2:: scc2L1277C_HIS6_FLAG6* | This study | B3042 |
|  | *MATa, SCC1-PK9::KANMX, scc2::NATMX4*  *smc1E1102Cmyc9:: kiTRP1, Δlys2:: scc2T1281C_HIS6_FLAG6* | This study | B3043 |
|  | *MATa, SCC1-PK9::KANMX, scc2::NATMX4*  *smc1E1102Cmyc9:: kiTRP11, Δlys2:: scc2R1289C_HIS6_FLAG6* | This study | B3037 |
|  | *MATa, SCC1-PK9::KANMX, scc2::NATMX4*  *smc1E1102Cmyc9:: kiTRP1, Δlys2:: scc2L1302C_HIS6_FLAG6* | This study | B3037 |
|  | *MATa, SCC1-PK9::KANMX, scc2::NATMX4*  *smc1E1102Cmyc9:: kiTRP1, Δlys2:: scc2L1307C_HIS6_FLAG6* | This study | B3038 |
|  | *MATa, SCC1-PK9::KANMX, scc2::NATMX4*  *smc1E1102Cmyc9:: kiTRP1, Δlys2:: scc2L1315C_HIS6_FLAG6* | This study | B3040 |
|  | *MATa, SCC1-PK9::KANMX, scc2::NATMX4*  *smc1E1102Cmyc9:: kiTRP1, Δlys2:: scc2L1325C_HIS6_FLAG6* | This study | B3041 |
|  | *MATa, SCC1-PK9::KANMX, scc2::NATMX4*  *smc1E1102Cmyc9:: kiTRP1, Δlys2:: scc2Y1279C_HIS6_FLAG6* | This study | B3074 |
|  | *MATa, SCC1-PK9::KANMX, scc2::NATMX4*  *smc1E1102Cmyc9:: kiTRP1, Δlys2:: scc2L1280C_HIS6_FLAG6* | This study | B3075 |
|  | *MATa, SCC1-PK9::KANMX, scc2::NATMX4*  *smc1E1102Cmyc9:: kiTRP1, Δlys2:: scc2V1283C_HIS6_FLAG6* | This study | B3076 |
|  | *MATa, SCC1-PK9::KANMX, scc2::NATMX4*  *smc1E1102Cmyc9:: kiTRP1, Δlys2:: scc2S1285C_HIS6_FLAG6* | This study | B3077 |
|  | *MATa, SCC1-PK9::KANMX, scc2::NATMX4*  *smc1E1102Cmyc9:: kiTRP1, Δlys2:: scc2L1287C_HIS6_FLAG6* | This study | B3078 |
|  | **Figure 4 and S4** |  |  |
| 4B and S4E | *Mat a,Δsmc3::HIS3, SCC1-PK9::KANMX, Δmet15::smc3M74C_HA3, Δscc2::NATMX4*  *Δlys2:: scc2N780C_HIS6_FLAG6* | This study | B3274 |
|  | *Mat a, Δsmc3::HIS3, SCC1-PK9::KANMX*  *Δmet15::smc3M74C_HA3, Δscc2::NATMX4*  *Δlys2:: scc2D782C_HIS6_FLAG6* | This study | B3276 |
|  | *Mat a, Δsmc3::HIS3, Δmet15::smc3M74C_HA3, SCC1-PK9::KANMX, Δscc2::NATMX4, Δlys2:: scc2*  *E819C_HIS6_FLAG6* | This study | B3272 |
|  | *Mat a, SCC1-PK9::KANMX, Δsmc3::HIS3, Δmet15::sm3M74C_HA3, Δscc2::NATMX4*  *Δlys2:: scc2D820C_HIS6_FLAG6* | This study | B3268 |
|  | *Mat a, SCC1-PK9::KANMX, Δsmc3::HIS3, Δmet15::smc3M74C_HA3, Δscc2::NATMX4*  *Δlys2:: scc2E821C_HIS6_FLAG6* | This study | B3323 |
|  | *Mat a, Δscc2::NATMX4, Δsmc3::HIS3*  *SCC1-PK9::KANMX, Δmet15::smc3M74C_HA3*  *Δlys2::Scc2R830C_HIS6_FLAG6* | This study | B3432 |
|  | *Mat a, Δscc2::NATMX4, Δsmc3::HIS3, SCC1-PK9::KANMX*  *Δmet15::smc3M74C_HA3, Δlys2:: scc2L878C_HIS6_FLAG6* | This study | B3433 |
|  | *Mat a, Δscc2::NATMX4, Δsmc3::HIS3, SCC1-PK9::KANMX*  *Δmet15::smc3M74C_HA3, Δlys2:: scc2N886C_HIS6_FLAG6* | This study | B3434 |
| 4C | *Mat a, smc3E202C-HA6::spHis5, SCC1-PK9:: NATMX, SCC2_6xHis_FLAG6::KANMX,pBH1194 (smc1(R1031C_E1102TAG)-myc), pBH61 (tRNATyr/Bpa synthetase)* | This study | B3227 |
|  | *Mat a, SCC1-PK9::KANMX, Δsmc1::NATMX4, Δmet15::smc1R1031C_Myc9, SCC2_6xHis_FLAG6::KANMX, pBH1092(smc3E202C_Q67TAG-HA3), pBH61 (tRNATyr/Bpa synthetase)* | This study | B2285 |
| 4D | *Mat a, SCC1-PK9::KANMX, Δsmc1::NATMX4, Δmet15::smc1N1192C_Myc9, SCC2_6xHis_FLAG6::KANMX, pBH1092(smc3R1222C_Q67TAG-HA3),pBH61 (tRNATyr/Bpa synthetase)* | This study | B3009 |
|  | *Mat a, SCC1-PK9::NATMX, smc3R1222C_HA6::SpHis3*  *SCC2_6xHis_FLAG6::KANMX, pBH1121 (smc1N1192C_E1102TAG-myc9), pBH61 (tRNATyr/Bpa synthetase)* | This study | B4041 |
| S4G | *MAT a, Δsmc1::NATMX4, SCC1-PK9::KANMX Δmet15::smc1K191CMyc9,Δsmc3::HIS3, ura3::smc3K184C-HA3 (pBH986)::URA3* | This study | B2656 |
|  | *Mat alpha, Δsmc1::NATMX4, SCC1-PK9::KANMX*  *Δmet15::smc1K191CMyc9 , Δsmc3::HIS3*  *ura3::smc3E188C-HA3::URA3* | This study | B2612 |
|  | *MAT a, SCC1-PK9::KANMX smc3Q195C-HA6::His3MX6*  *Δsmc1::NATMX4, leu2::smc1S195C-myc9::HPHNT1* | This study | B2423 |
|  | *Mat a,*  *SCC1-PK9::KANMX Δsmc1::NATMX4*  *leu2::smc1K201C-myc9::HPHNT1*  *smc3K198C-HA6::His3MX6* | This study | B2395 |
|  | *Mat a,*  *Δsmc1::NATMX4, Smc3E202C-HA6::spHis5*  *SCC1-PK9::KANMX Δmet15:sSmc1E209C-myc9* | This study | B2311 |
|  | *Mat a,*  *Δsmc1::NATMX4, smc3E202C-HA6::spHis5*  *SCC1-PK9::KANMX Δmet15::smc1R1031C-myc9* | This study | B2233 |
|  | *Mat a,*  *Δsmc1::NATMX4, SCC1-PK9::KANMX Δmet15::smc1R1020C-myc9*  *smc3E213C-HA6::His3MX6* | This study | B2376 |
|  | **Figure 5 and S5** |  |  |
| 5A and S5A | *MATa, SCC1-PK9::KANMX, Δscc2::NATMX4*  *Δlys2:: scc2T1175C_HIS6_FLAG6, Δmet15::smc3Q67C-HA3, leu2::GAL1P-SIC1(9m)/HIS3PGAL1/HIS3P-GAL2/GAL1P-GAL4::LEU2* | This study | B3262 |
|  | *MATa, SCC1-PK9::KANMX, Δscc2::NATMX4*  *Δlys2:: scc2T1175C_HIS6_FLAG6, Δmet15:: smc3(K112Q_K113Q_Q67C) -HA3, leu2::GAL1P-SIC1(9m)/HIS3PGAL1/HIS3P-GAL2/GAL1P-GAL4::LEU2* | This study | B3263 |
| 5D and S5C | *Mat a, SCC1-PK9::NATMX, Scc2_6xHis_FLAG6::KANMX*  *Δsmc3::HIS3, Δmet15::Smc3_HA3* | This study | B3476 |
|  | *Mat a, SCC1-PK9::NATMX, scc2E819C_HIS6_FLAG6::KANMX, Δsmc3::HIS3, Δmet15::Smc3_HA3* | This study | B3475 |
|  | *Mat a, Δsmc3::HIS3, SCC1-PK9::NATMX, Δmet15::smc3S72C_HA3, SCC2_6xHis_FLAG6::KANMX* | This study | B3473 |
|  | *Mat a, Δsmc3::HIS3, SCC1-PK9::NATMX*  *Δmet15::smc3S72C_HA3, scc2E819C_HIS6_FLAG6::KANMX* | This study | B3471 |
| 5E and S5D | *MATa,SCC1-PK9::NATMX, SMC3HA6::HIS3*  *SCC2_6xHis_FLAG6::KANMX, leu2::GAL1P-SIC1(9m)/HIS3PGAL1/HIS3P-GAL2/GAL1P-GAL4::LEU2* | This study | B3913 |
|  | *MATa, SCC1-PK9::NATMX*  *scc2_D369C_FLAG6::KanMX, SMC3-HA6::HIS3*  *leu2::GAL1P-SIC1(9m)/HIS3PGAL1/HIS3P-GAL2/GAL1P-GAL4::LEU2* | This study | B3929 |
|  | *MATa,SCC1-PK9::NATMX, smc3_K1004C_HA6::His3*  *SCC2_6xHis_FLAG6::KANMX, leu2::GAL1P-SIC1(9m)/HIS3PGAL1/HIS3P-GAL2/GAL1P-GAL4::LEU2* | This study | B3927 |
|  | *MATa, SCC1-PK9::NATMX, Smc3_K1004C_HA6::His3*  *scc2_D369C_FLAG6::KanMX, leu2::GAL1P-SIC1(9m)/HIS3PGAL1/HIS3P-GAL2/GAL1P-GAL4::LEU2* | This study | B3916 |
| 5F and S5E | *Mat a, SCC1-PK9::KANMX, Δmet15:smc3S72C_HA3, Δscc2::NATMX4, Δlys2::scc2E819C_HIS6_FLAG6*  *leu2::GAL1P-SIC1(9m)/HIS3PGAL1/HIS3P-GAL2/GAL1P-GAL4::LEU2* | This study | B3371 |
|  | *Mat a, SCC1-PK9::KANMX, Δmet15::smc3(K112Q_K113Q_S72C)-HA3, Δscc2::NATMX4, Δlys2::scc2E819C_HIS6_FLAG6*  *leu2::GAL1P-SIC1(9m)/HIS3PGAL1/HIS3P-GAL2/GAL1P-GAL4::LEU2* | This study | B3417 |
| 5H and S5F | *MATa, SCC1-PK6::TRP1, Ura3::smc3_K1004C_HA3::URA3, scc2_D369C_FLAG6::KanMx, leu2::GAL1P-SIC1(9m)/HIS3PGAL1/HIS3P-GAL2/GAL1P-GAL4::LEU2* | This study | B3814 |
|  | *MATa,SCC1-PK6::TRP1, Ura3::smc3_( K112Q_K113Q_K1004C)-HA3::URA3, scc2_D369C_FLAG6::KanMx, leu2::GAL1P-SIC1(9m)/HIS3PGAL1/HIS3P-GAL2/GAL1P-GAL4::LEU2* | This study | B3815 |
| 5J | *MATa, Δscc2::NATMX4, smc1E1102Cmyc9:: kiTRP1*  *Δlys2:: scc2T1281C_HIS6_FLAG6, ura3::smc3(K112Q, K113Q)-PK6::URA3 , leu2::GAL1P-SIC1(9m)/HIS3PGAL1/HIS3P-GAL2/GAL1P-GAL4::LEU2* | This study | B3265 |
|  | *MATa, Δscc2::NATMX4, smc1E1102Cmyc9:: kiTRP1, Δlys2:: scc2T1281C_HIS6_FLAG6, ura3::SMC3-PK6::URA3 , leu2::GAL1P-SIC1(9m)/HIS3PGAL1/HIS3P-GAL2/GAL1P-GAL4::LEU2* | This study | B3266 |
|  | *MATa, Δscc2::NATMX4, smc1E1102Cmyc9:: kiTRP1, Δlys2:: scc2T1281C_HIS6_FLAG6, leu2::GAL1P-SIC1(9m)/HIS3PGAL1/HIS3P-GAL2/GAL1P-GAL4::LEU2* | This study | B3267 |
| S5B | *MATa, SCC1-PK9::KANMX, SCC2_6xHis_FLAG6::KANMX, pBH585 (smc3-Q67TAG-HA3), pBH61 (tRNATyr/Bpa synthetase)* | This study | B1505 |
|  | *MATa, SCC1-PK9::KANMX, SCC2_6xHis_FLAG6::KANMX, pBH323 (smc3QQ-Q67TAG-HA3), pBH61 (tRNATyr/Bpa synthetase)* | This study | B2762 |
|  | **Figure 6 and S6** |  |  |
| 6A and S6A | *MATa, , SCC1-PK9::KANMX, Δscc2::NATMX4, Δlys2::scc2T1175C_HIS6_FLAG6, Δmet15::smc3Q67C-HA3, leu2::GAL1P-SIC1(9m)/HIS3PGAL1/HIS3P-GAL2/GAL1P-GAL4::LEU2* | This study | B3262 |
|  | *MATa, , SCC1-PK9::KANMX, Δscc2::NATMX4, Δlys2:scc2T1175C_HIS6_FLAG6, Δmet15::smc3(K112Q_K113Q_Q67C)-HA3, leu2::GAL1P-SIC1(9m)/HIS3PGAL1/HIS3P-GAL2/GAL1P-GAL4::LEU2* | This study | B3263 |
|  | *MATa, , SCC1-PK9::KANMX, Δscc2::NATMX4, Δlys2:scc2T1175C_HIS6_FLAG6, Δmet15::smc3(K112Q_K113Q_Q67C-R1008I)-HA3, leu2::GAL1P-SIC1(9m)/HIS3PGAL1/HIS3P-GAL2/GAL1P-GAL4::LEU2* | This study | B3362 |
|  | *MATa, , SCC1-PK9::KANMX, Δscc2::NATMX4, Δlys2:scc2T1175C_HIS6_FLAG6, Δmet15::smc3(K112Q_K113Q_Q67C-W483R)-HA3, leu2::GAL1P-SIC1(9m)/HIS3PGAL1/HIS3P-GAL2/GAL1P-GAL4::LEU2* | This study | B3427 |
|  | *MATa, , SCC1-PK9::KANMX, Δscc2::NATMX4, Δlys2:scc2T1175C_HIS6_FLAG6, Δmet15::smc3(K112Q_K113Q_Q67C_R1008I_W483R)-HA3, leu2::GAL1P-SIC1(9m)/HIS3PGAL1/HIS3P-GAL2/GAL1P-GAL4::LEU2* | This study | B3363 |
| 6C and S6C | *Mat a, SCC1-PK6::TRP1, Ura3::smc3K1004C_HA3::URA3, scc2_D369C_FLAG6::KanMx, leu2::GAL1P-SIC1(9m)/HIS3PGAL1/HIS3P-GAL2/GAL1P-GAL4::LEU2* | This study | B3814 |
|  | *Mat a, SCC1-PK6::TRP1, Ura3::smc3K112Q_K113Q_K1004C_HA3::URA3, scc2_ D369C_FLAG6::KanMx, leu2::GAL1P-SIC1(9m)/HIS3PGAL1/HIS3P-GAL2/GAL1P-GAL4::LEU2* | This study | B3815 |
|  | *Mat a, SCC1-PK6::TRP1, Ura3::smc3K112Q_K113Q_K1004C_R1008I-HA3::URA3, scc2_ D369C_FLAG6::KanMx, leu2::GAL1P-SIC1(9m)/HIS3PGAL1/HIS3P-GAL2/GAL1P-GAL4::LEU2* | This study | B3816 |
|  | *Mat a, SCC1-PK6::TRP1, Ura3::smc3K112Q_K113Q_K1004C_W483R-HA3::URA3, scc2_ D369C_FLAG6::KanMx, leu2::GAL1P-SIC1(9m)/HIS3PGAL1/HIS3P-GAL2/GAL1P-GAL4::LEU2* | This study | B3817 |
|  | *Mat a, SCC1-PK6::TRP1, Ura3::smc3K112Q_K113Q_K1004C_W483R_R1008I-HA3::URA3, scc2_D369C_FLAG6::KanMx, leu2::GAL1P-SIC1(9m)/HIS3PGAL1/HIS3P-GAL2/GAL1P-GAL4::LEU2* | This study | B3818 |
| 6F | *MAT a/alpha, Δsmc3::HIS3/SMC3, ura3/ura3::smc3(K112Q_K113Q_R1008I)::URA3, scc2(E822K, L937F):: NATMX / SCC2* | This study | B1392 |
| 6G | *Mat a, ura3::SMC3-PK6::URA3* | This study | B4002 |
|  | *Mat a, ura3::smc3(K112Q_K113Q)-PK6:: URA3* | This study | B4003 |
|  | *Mat a, ura3::smc3(K112Q_K113Q_R1008I)-PK6:: URA3* | This study | B3945 |
|  | *Mat a, ura3::smc3-(K112Q_K113Q)-PK6:: URA3 ,Scc2(E822K, L937F):: NatMX* | This study | B4005 |
|  | *Mat a, ura3::smc3(K112Q_K113Q_R1008I)-PK6:: URA3, Scc2(E822K, L937F):: NatMX* | This study | B4006 |
| S6B | *MATa, SCC1-PK9::KANMX, SCC2_6xHis_FLAG6::KANMX, pBH585 (smc3-Q67TAG-HA3 ), pBH61 (tRNATyr/Bpa synthetase)* | This study | B1505 |
|  | *MATa, SCC1-PK9::KANMX,, SCC2_6xHis_FLAG6::KANMX, pBH623 (smc3QQ -Q67TAG-HA3 ), pBH61 (tRNATyr/Bpa synthetase)* | This study | B2762 |
|  | *MATa, SCC1-PK9::KANMX, SCC2_6xHis_FLAG6::KANMX pBH995(smc3QQ -Q67TAG-R1008I-HA3 ), pBH61 (tRNATyr/Bpa synthetase)* | This study | B2757 |
|  | *MATa, SCC1-PK9::KANMX, SCC2_6xHis_FLAG6::KANMX, pBH998 (smc3-Q67TAG-W483R-HA3 ), pBH61 (tRNATyr/Bpa synthetase)* | This study | B2760 |
|  | *MATa, SCC1-PK9::KANMX, SCC2_6xHis_FLAG6::KANMX, pBH999 (smc3-Q67TAG-W483R-R1008I-HA3 ), pBH61 (tRNATyr/Bpa synthetase)* | This study | B2761 |
| S6D | *Mat a, SCC1-PK9::KANMX, Δmet15::smc3S72C_HA3, Δscc2::NATMX4, Δlys2::scc2E819C_HIS6_FLAG6*  *leu2::GAL1P-SIC1(9m)/HIS3PGAL1/HIS3P-GAL2/GAL1P-GAL4::LEU2* | This study | B3371 |
|  | *Mat a, SCC1-PK9::KANMX, Δmet15::smc3(K112Q_K113Q_S72C)-HA3, Δscc2::NATMX4, Δlys2::scc2E819C_HIS6_FLAG6*  *leu2::GAL1P-SIC1(9m)/HIS3PGAL1/HIS3P-GAL2/GAL1P-GAL4::LEU2* | This study | B3417 |
|  | *Mat a, SCC1-PK9::KANMX, Δmet15::smc3(K112Q_K113Q_S72C_R1008I)-HA3, Δscc2::NATMX4, Δlys2::scc2E819C_HIS6_FLAG6*  *leu2::GAL1P-SIC1(9m)/HIS3PGAL1/HIS3P-GAL2/GAL1P-GAL4::LEU2* | This study | B3593 |
|  | *Mat a, SCC1-PK9::KANMX, Δmet15::smc3(K112Q_K113Q_S72C_W483R)-HA3, Δscc2::NATMX4, Δlys2::scc2E819C_HIS6_FLAG6*  *leu2::GAL1P-SIC1(9m)/HIS3PGAL1/HIS3P-GAL2/GAL1P-GAL4::LEU2* | This study | B3589 |
|  | *Mat a, SCC1-PK9::KANMX, Δmet15::smc3(K112Q_K113Q_S72C_W483R_R1008I)-HA3, Δscc2::NATMX4, Δlys2::scc2E819C_HIS6_FLAG6*  *leu2::GAL1P-SIC1(9m)/HIS3PGAL1/HIS3P-GAL2/GAL1P-GAL4::LEU2* | This study | B3591 |
| S6G | *MATa/alpha, SMC3/Δsmc3::HIS3*  *ura3/ura3::Smc3(K112Q_K113Q_R1008I)::URA3*  *SCC2/Scc2(E822K):: NATMX* | This study | B3206 |
|  | *MATa/alpha, SMC3/Δsmc3::HIS3 ura3/ura3::Smc3(K112Q_K113Q_R1008I)::URA3*  *SCC2/scc2(L937F):: NATMX* | This study | B3207 |
|  | **Figure 7 and S7** |  |  |
| 7A and S7 | *Mat a, SCC1-PK6::TRP1, ura3::smc3Q67C-HA3::URA3, Scc2T1175C_HIS6_FLAG6::KANMX, leu2::GAL1P-SIC1(9m)/HIS3PGAL1/HIS3P-GAL2/GAL1P-GAL4::LEU2* | This study | B3819 |
|  | *Mat a, SCC1-PK6::TRP1, ura3::smc3Q67C_E1155Q-HA3::URA3, scc2T1175C_HIS6_FLAG6::KANMX*  *leu2::GAL1P-SIC1(9m)/HIS3PGAL1/HIS3P-GAL2/GAL1P-GAL4::LEU2* | This study | B3820 |
|  | *MATa, SCC1-PK9::NATMX, ura3::smc3_S72C_HA3::URA3*  *scc2E819C_HIS6_FLAG6::KANMX, leu2::GAL1P-SIC1(9m)/HIS3PGAL1/HIS3P-GAL2/GAL1P-GAL4::LEU2* | This study | B3821 |
|  | *MATa, SCC1-PK9::NATMX, ura3::smc3_S72C_E1155Q_HA3::URA3, scc2E819C_HIS6_FLAG6::KANMX, leu2::GAL1P-SIC1(9m)/HIS3PGAL1/HIS3P-GAL2/GAL1P-GAL4::LEU2* | This study | B3822 |
|  | *MATa, SCC1-PK9::NATMX*  *ura3::smc3_K1004C_HA3::URA3, scc2E819C_HIS6_FLAG6::KANMX, leu2::GAL1P-SIC1(9m)/HIS3PGAL1/HIS3P-GAL2/GAL1P-GAL4::LEU2* | This study | B3880 |
|  | *MATa, SCC1-PK9::NATMX, ura3::smc3_K1004C_E1155Q_HA3::URA3, scc2E819C_HIS6_FLAG6::KANMX, leu2::GAL1P-SIC1(9m)/HIS3PGAL1/HIS3P-GAL2/GAL1P-GAL4::LEU2* | This study | B3890 |
|  | *MATa, SCC1-PK9::NATMX*  *ura3::smc1E1102C-MYC9::URA3*  *scc2T1281C_HIS6_FLAG6::KANMX, leu2::GAL1P-SIC1(9m)/HIS3PGAL1/HIS3P-GAL2/GAL1P-GAL4::LEU2* | This study | B3823 |
|  | *MATa, SCC1-PK9::NATMX, ura3::smc1E1102C_E1158Q-MYC9::URA3, scc2T1281C_HIS6_FLAG6::KANMX*  *leu2::GAL1P-SIC1(9m)/HIS3PGAL1/HIS3P-GAL2/GAL1P-GAL4::LEU2* | This study | B3824 |
| 7C | *MATa, SCC1-PK9::HYGMX, Ura3::smc3_S72C_K1004C-HA3::URA3, scc2D369C_E819C_HIS6_FLAG6::KANMX*  *leu2::GAL1P-SIC1(9m)/HIS3PGAL1/HIS3P-GAL2/GAL1P-GAL4::LEU2* | This study | B3892 |
|  | *MATa, SCC1-PK9::HYGMX, Ura3::smc3_K1004C_E1155Q-HA3::URA3, scc2D369C_HIS6_FLAG6::KANMX*  *leu2::GAL1P-SIC1(9m)/HIS3PGAL1/HIS3P-GAL2/GAL1P-GAL4::LEU2* | This study | B3890 |
|  | *MATa, SCC1-PK9::HYGMX, Ura3::smc3_S72C_E1155Q-HA3::URA3, scc2D369C_E819C_HIS6_FLAG6::KANMX*  *leu2::GAL1P-SIC1(9m)/HIS3PGAL1/HIS3P-GAL2/GAL1P-GAL4::LEU2* | This study | B3822 |
|  | *MATa, SCC1-PK9::HYGMX, Ura3::smc3_S72C_K1004C-E1155Q-HA3::URA3, scc2D369C_E819C_HIS6_FLAG6::KANMX*  *leu2::GAL1P-SIC1(9m)/HIS3PGAL1/HIS3P-GAL2/GAL1P-GAL4::LEU2* | This study | B3891 |
